# *Viola unica* (Violaceae L.), a very rare, high elevation, single-site new species endemic to the northern semi-desert Altiplano of Chile

**DOI:** 10.1101/787564

**Authors:** John Michael Watson, Ana Rosa Flores

## Abstract

While examining the *Viola* collection at the herbarium (SGO) of the Museo Nacional de Historia Natural (MNHN), Santiago, we encountered a folder with a single unidentified specimen consisting of a solitary rosette. It had been found in northern Chile near the border with Bolivia, where considerable mining activity takes place. The plant appeared to differ from all others related to it as known to ourselves, and on investigation it did indeed prove to be undescribed. It is presented herein, together with a detailed analysis and preliminary partial revision of related taxa, including a key.

## Introduction, *Viola* and its section *Andinium* W. Becker

*Viola* is cosmopolitan and of mainly temperate and high tropical mountain distribution. With an estimated 610-655 known and accepted species it is the largest genus of its family, Violaceae, and is comprised of 16 sections, the majority endemic to the Northern Hemisphere (Wahlert *et al*. 2014, Watson & Flores 2018b). *Viola* has been found by investigators to have evolved ca. 35 Ma years ago in what is now the southern end of temperate South America (Clausen 1929, Ballard *et al*. 1999, Marcussen *et al*. 2012, Marcussen *et al*. 2015). The location of this early branching from the rest of the family has led to the conclusion that species of the most direct ancestral origin exist in the subcontinent in three of its sections, two of which are endemic, the third being mainly distributed there.

The largest of those 16 sections, when its undescribed but accepted species are taken into account, is sect. *Andinium* W. Becker (Watson & Flores 2018b), one of the two endemic to South America. Its 108 published taxa (IPNI 2019, Watson *et al*. 2019, Watson & Flores 2019a, b), together with the present species and others waiting to be described or collected, are known coloquially as the Andean rosulate violas. With a longitudinal distribution between the equator and southern Patagonia, their full discovered compliment amounts to at least 147 species as currently recognised by the present authors. This total includes 39-40 known either as specimens - or in a few instances as reliable photographs - which are as yet unpublished (Watson & Flores ined.).

Apart effectively from two significant regional floras, sect. *Andinium* was ignored botanically for decades following the death in 1928 of its historical authority, Wilhelm Becker of Berlin-Dahlem, with specialised study of the infrageneric alliance as a whole not resumed until the mid-1990s. When these factors are taken in conjunction with the inaccessible habitats of many of its taxa, the majority as solitary or few known populations, the fact that it is still poorly understood compared with the other sections is hardly surprising. Furthermore, as many as 40 species are currently unknown in the wild, which serves to compound the problems, as also do the destruction of Becker’s large and important collection at Berlin-Dahlem during WW2 (Hiepko 1987, Haagemann & Zepernick 1993) and the difficulty of distinguishing between some taxa. Nevertheless, Marcussen *et al*. (2015) have been able to calculate that the section split from the rest of *Viola* as early as 29 Ma. This revelation, together with the section’s specialised adaptation to developing Andean uplift and more recent Mediterranean geoclimatic conditions, explains why so many of its taxa are uniquely unlike the rest of *Viola* (Watson & Flores 2012, 2013a, 2013b).

The novelty described below, as lodged in the SGO collection, is exemplary of the difficulties involved in providing accurate information on many sect. *Andinium* taxa. It is one of those currently unknown in the wild, its exact whereabouts being unrecorded, and with no collector to consult for details such as population size and whether it is has been found at more than one site. Nevertheless, its extreme rarity and situation in a sector where widescale copper mining is in progress, with further potential operations in view there, invoke the urgent need for its existence to be made known as far as is possible under the circumstances. This is the first urgent and vital step towards the aspiration that it may be located and effectively protected.

## Materials and methods

### *Viola* sect

*Andinium* has been the main focus of our botanical investigation for 25 years. We are therefore familiar with all its taxa as described in the literature, in addition to many observed as specimens during exhaustive examinations of collections of the section at B, CONC, K, LIL, MERL, P, SGO, SI and ULS, together with those we and others have encountered in the field. This enabled us to recognise with confidence that the taxon herein has not hitherto been described, and also to appreciate how it relates to and differs from those in the section we judge to be most closely related to it.

The specimen was soaked and observed under high magnification, when details not readily evident to the naked eye were noted, and accurate measurements taken as well as photographs.

### Taxonomic result

***Viola unica*** J.M. Watson & A.R. Flores, sp. nov. (Figs. 1-4)

**Fig. 1.**
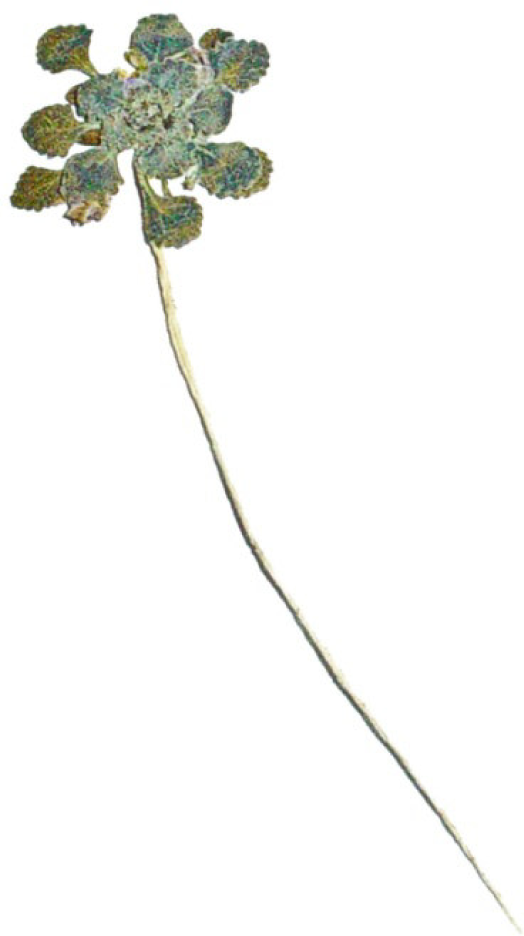
The pressed herbarium type specimen of *Viola unica*. (Photo - A.R. Flores, 25 Aug 2008)

**Fig. 2.**
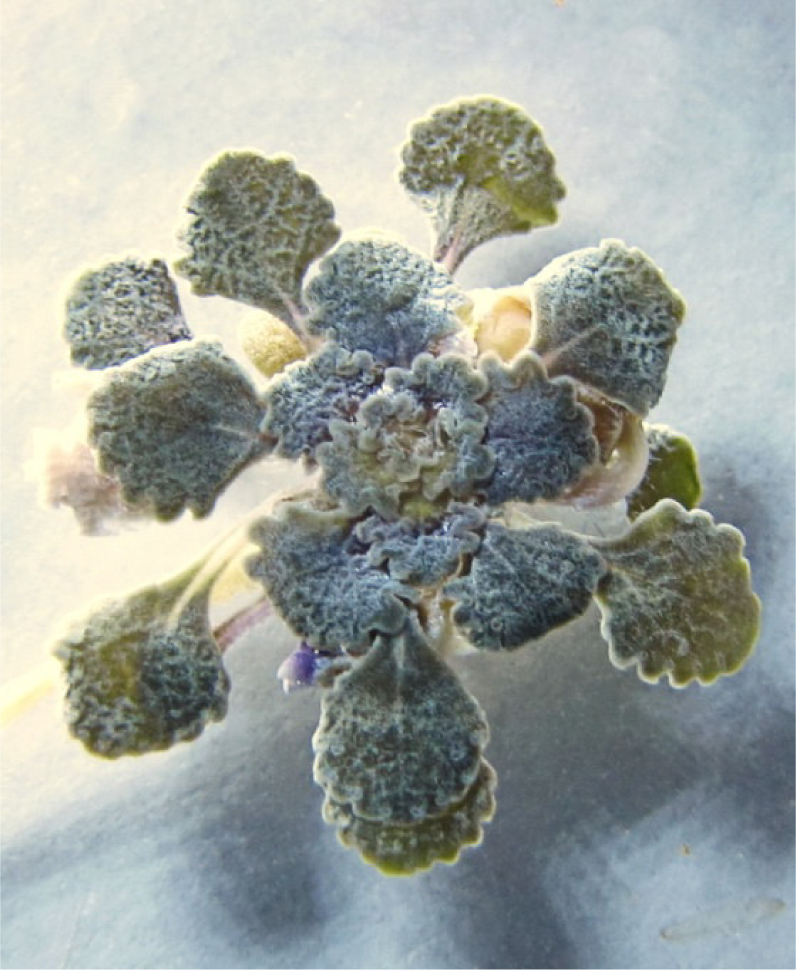
Plan view of rosette of soakedtype specimen of *Viola unica*. (Photo - A.R. Flores, 7 Jun 2019)

**Fig. 3.**
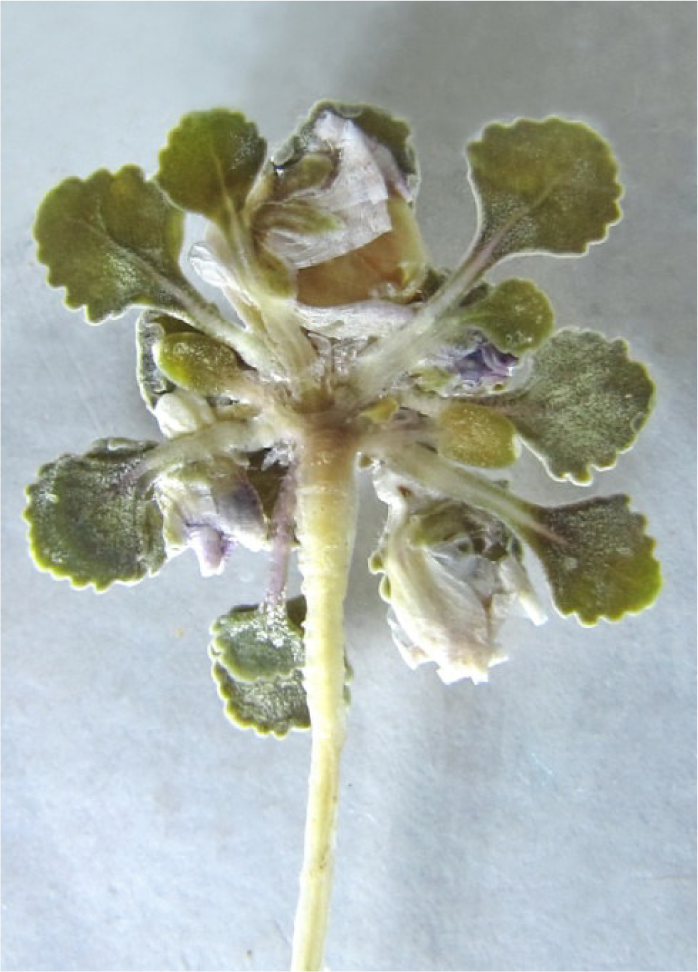
Underside of rosette of soaked type specimen of *Viola unica*. (Photo - A.R. Flores, 7 June 2019)

**Fig. 4.**
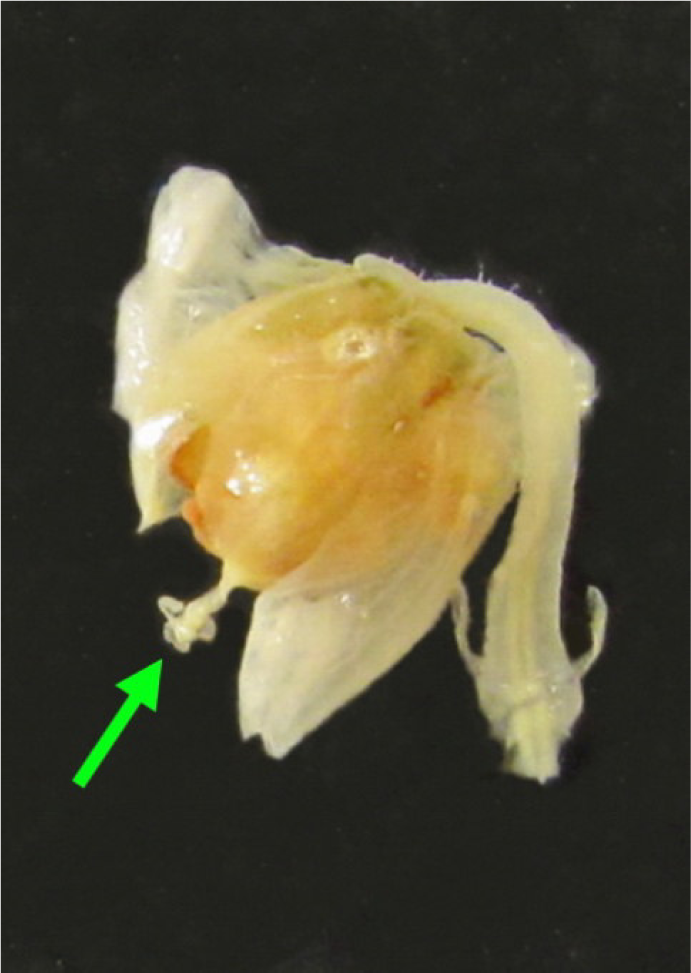
Preserved *Viola unica* flower showing peduncle with basal bracteoles, ripe capsule, and attached trilobed style crest arrowed green. (Photo - A.R. Flores, 7 Jun 2019)

#### Type

CHILE. Tarapacá Region, Tamarugal Province, Pica Community near Collahuasi, approximate coordinates 21°00’S 68°00’W, elevación ca. 4000-4500 m, Feb 2008, collector unknown, specimen number P15 (holotype SGO).

#### Diagnosis

The new species is disinct from all known others of sect. *Andinium* W. Becker. Among the perennials, its discretely trilobed style crest is a morphological feature shared with the four described taxa of the informal *Triflabellatae* W. Becker alliance and *Viola flos-idae* Hieron., which differs from *V. unica* by the glandular, not eglandular undersurface of its laminas. *Viola mesadensis* W. Becker possesses apical and lateral crest appendages, but also differs from *V. unica* by its glandular lamina undersurface as well as its glabrous, not bearded lateral petals. Except for *V. mesadensis* and at times *V. flos-idae*, all those taxa have a narrowly elliptical to ovate lamina with an acute apex as opposed to the rounded, obtuse blades of *V. unica*. The latter foliar morphology as well as the species’ regularly crenate leaves ally it more with several species related to *Viola volcanica* Gillies ex Hook. & Arn.; also with *Viola beati* J.M. Watson & A.R. Flores and *Viola singularis* J.M. Watson & A.R. Flores. But the style crests are apical for the first named and lateral only for the following two. In addition the strongly curved lobes are unique, being unknown in any other trilobed species of the section, except for one very distantly related annual.

#### Description

*Life form* perennial (or annual, see notes 1 and 3 below), acaulous, rosulate, evergreen hemicryptophyte. *Rootstock* axial, narrowly flagelliform, fibrous, ca. 7 cm long × 1 mm dia. at junction with caudex. *Caudex* 3 mm, naked (on specimen, but see notes 1 and 3 below). *Plant* (as known) solitary with single *rosette*, this 19 mm dia., open-structured, plane. *Leaves* spathulate, 7-12 mm when mature; *pseudopetioles* 4-7.5 × 0.5-0.7 mm, plane, fleshy; *stipules* 1.3 × 0.5 mm, triangular, hyaline, apex blunt; *lamina* 3-3.5 × 3-4.5 mm, rotund-ovate, approximately as wide as long or somewhat wider, narrowly cuneate to pseudopetiole, 4-5 rounded-crenate on each lateral margin, somewhat carnose, face dark, dull greyish green, alveolate reticulate, undersurface dark greyish, minutely farinose-papillose, margins usually glabrous, occasionally few-ciliate at base, apex rounded to shallow subtruncate-undulate. *Flowers* axial, solitary, situated within circumference of rosette. *Peduncles* to ca. 9 mm, shorter than mature leaves, sparsely pilose apically; *bracteoles* adnate with base of peduncle for 0.2 mm, free and spreading above, when 1.5 mm long × 0.6 mm wide, narrowly ovate, hyaline, apex acute. *Sepals* free, entire, triangular-lanceolate, subacute, herbaceous with broad hyaline margin; *superior sepal* 1.5-2 × 1-1.2 mm; lateral sepals 1.7-2.2 × 1-1.2 mm; inferior sepals 2-2.2 × 1-1.2 mm. *Corolla* white or pale, reverse of petals stained diffuse violet at base (see also note 2 below); *superior petals* 5 × 2-2.1 mm, oblanceolate, apex rounded; *lateral petals* 5.5 × 2.5 mm, upcurved-oblanceolate, apex rounded, upper half of face except apex bearded with short, stout, hyaline, clavate indumentum, as also present and suberect on adjacent upper margin; *inferior petal* 6 × 6.5 mm, broadly obcordate, sides incurved towards base, apical sinus broad, evenly curved; apical lobes rounded; *spur* 2 mm long × 1.5 mm dia., cylindrical, apex bluntly rounded. *Anthers* ca. 1.2 × 0.8-1 mm, lower pair with 1.2 mm filiform nectar spurs; *connectives* slightly shorter than anthers, ca. 1 mm, dull yellow. *Style* 0.7 mm, straight, clavate; *stigma* small frontal aperture on style head; *style crest* as long, linear, down- and incurved lobe with acute apex, situated on either side of style head, and third lobe apical, obtusely obovate, porrectly directed, with downcurved tip. *Fruit* ca. 5 mm, orbicular, tri-valved, capsule; *seeds* not seen.

#### Note 1

The life duration of the new species is difficult to assess from the solitary specimen with a single rosette and without remnants of any previous seasons’ growth on the caudex. In conjunction with the equivalent longevity of most allied species, the main evidence favouring a perennial life-form is the length of the rootstock and its relative thickness at the crown-junction with the rosette. Also, evident live cotyledons are present at the base of the rosette, which suggests the plant germinated fairly recently, is not yet mature, and was therefore likely to develop significantly. (Figs. 1-3)

#### Note 2

Floral pigmentation is eliminated or changed by pressing and drying, and the only way of knowing the exact living colour of the corolla is from accurate notes made by the collector, photographs (neither in this case), or knowledge of the plant *in vivo*. However, it is reasonable to speculate from the saturated specimen that the internal ground colour is white, very pale violet or lilac, probably with darker basal veining or dense streaking on the inferior and perhaps lateral petals, and certainly with a yellow throat to the inferior petal. This would be consistent with all other close relatives of the species.

#### Note 3

The flower is notably and atypically large relative to the foliage and rosette. This may be a standard feature of the species. Alternatively, the rosette may be an undeveloped immature perennial in its first season, as noted above, bearing regular sized flowers. Such morphology is known in four other species of the section with a southern distribution (Watson & Flores ined.) Alternatively, it may perhaps support the lesser possibility that *V. unica* is in fact annual.

#### Distribution

The new species is only known from one collection of a single individual found in the southern sector of Tarapacá Region in Chile. It is therefore a regional and national endemic respectively. (Figs. 5-6)

**Fig. 5.**
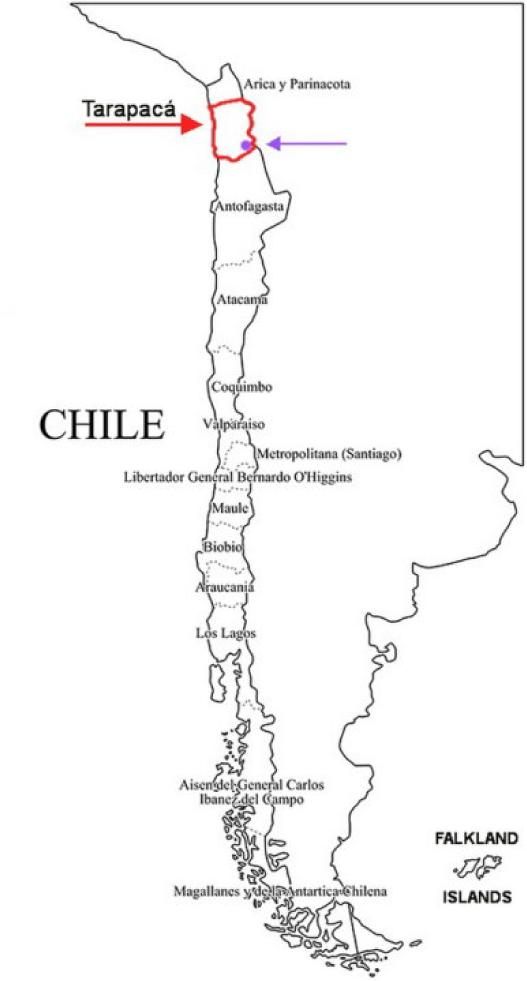
South America. The regions of Chile with Tarapacá, where *Viola unica* is endemic, outlined and arrowed red. General location of *Viola unica* arrowed violet.

**Fig. 6.**
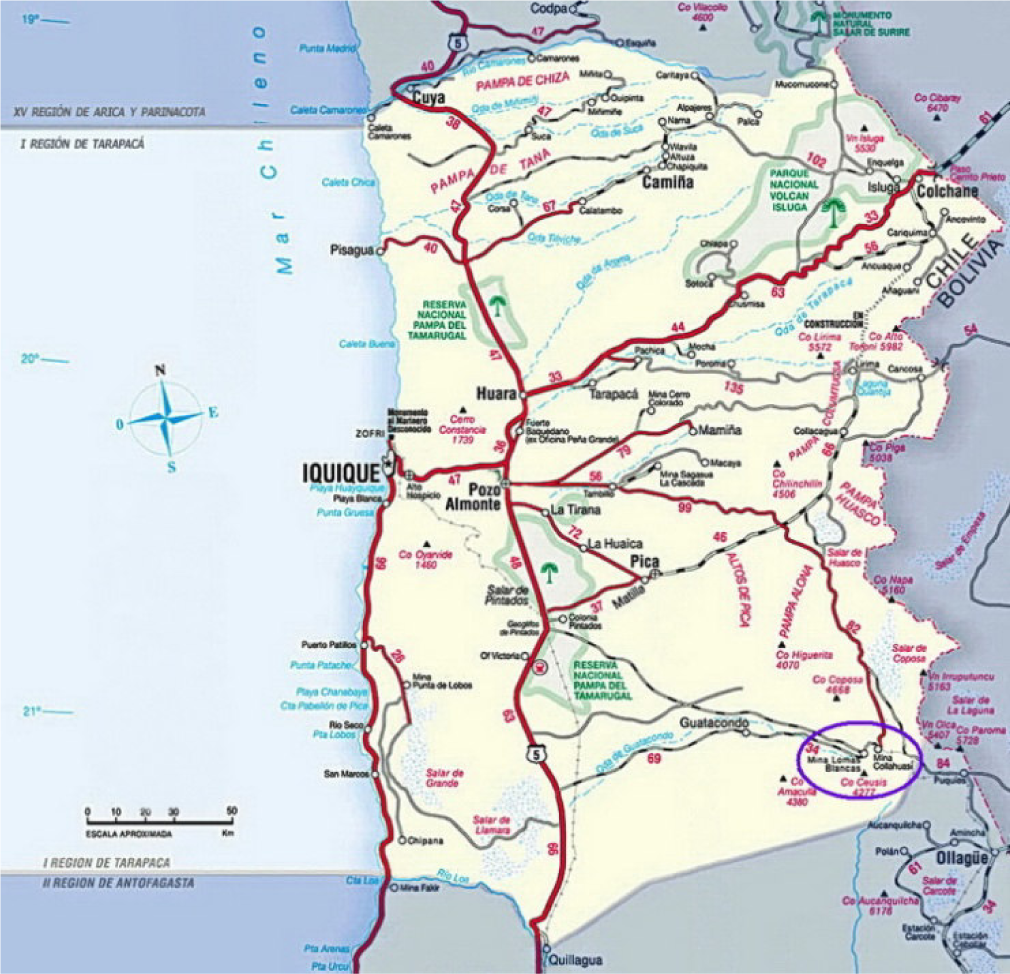
Tarapacá Region, northern Chile, with violet-ringed area within which *Viola unica* occurs. (Courtesy of Turistel)

#### Habitat

Gajardo (1994) classified the ecosystem where *V. unica* was found as ‘Estepa Alto-Andina Sub-Desértica’, which translates as ‘semi-desert high Andean steppe’. It is frequently domintated by extensive stretches of individual wiry bunch grasses (Poaceae) of the species *Anatherostipa venusta* (Phil.) Peñailillo and *Festuca chrysophylla* Phil. (fig. 7). Dispersed, compact, dense cushion and mat-forming species and dwarf xerophytic shrubs resistent to the effects of the extreme climate at high elevations characterize the petalloid flora. These include *Azorella compacta* Phil. (Apiaceae), *Baccharis tola* Phil. (Asteraceae), *Calceolaria stellariifolia* Phil. (Calceolariaceae) (fig. 8), *Chuquiraga spinosa* Less. (Asteraceae), *Fabiana bryioides* Phil. (Solanaceae) (fig. 9), *Jaborosa parviflora* (Phil.) Hunz. & Barbosa (Solanaceae) (fig. 10), *Lampayo medicinalis* Phil. (Verbenaceae), *Pycnophyllum molle* J. Rémy (Caryophyllaceae), and occasional examples of *Maihueniopsis boliviana* (Salm-Dyck) R. Kiesling (Cactaceae) (fig. 11).

**Fig. 7.**
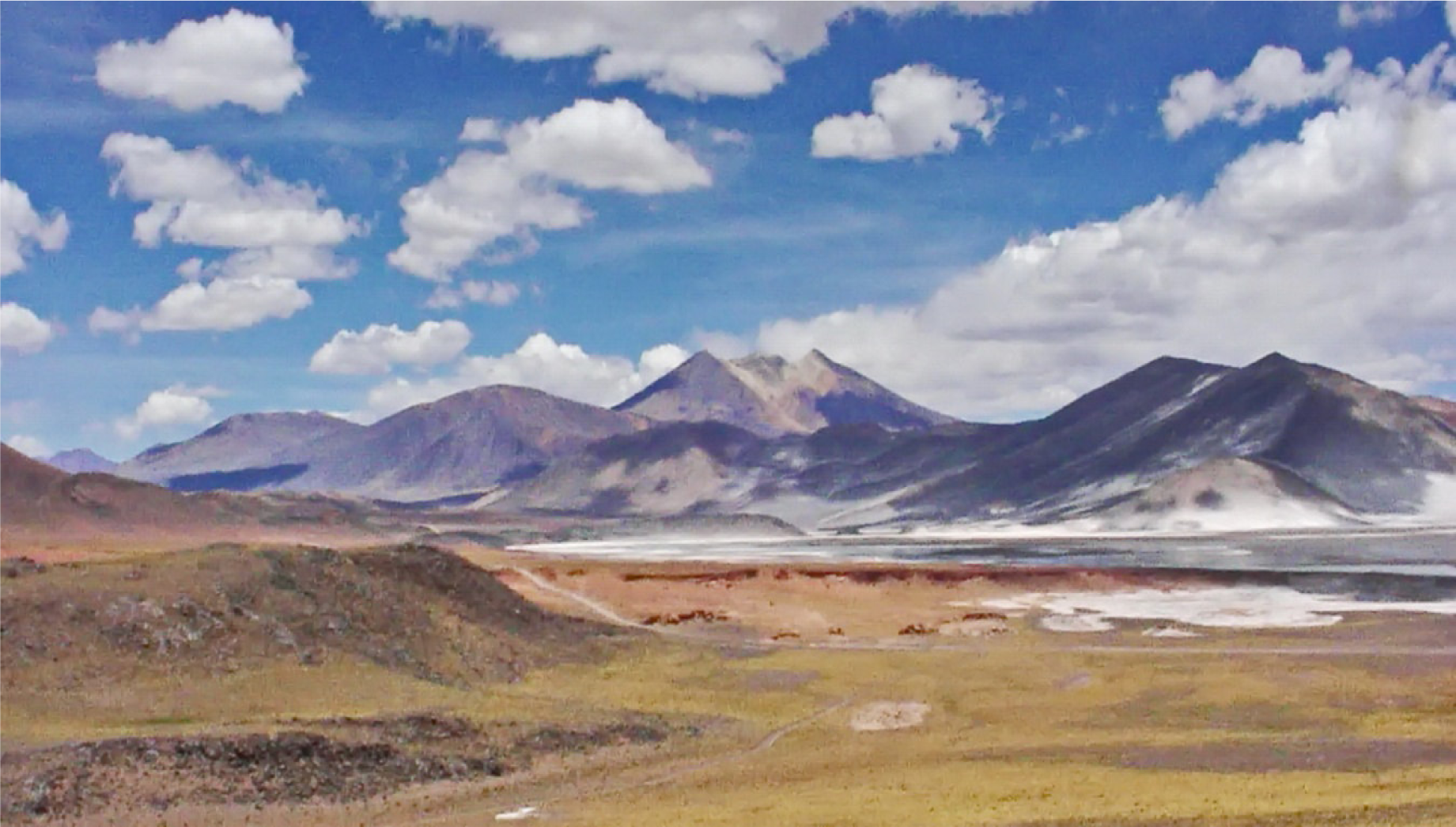
Typical Chilean high Altiplano landscape and vegetation, shortly to the south of Collahuasi, type area of *Viola unica*. (Photo - A.R. Flores, 6 Jan 2006)

**Fig. 8.**
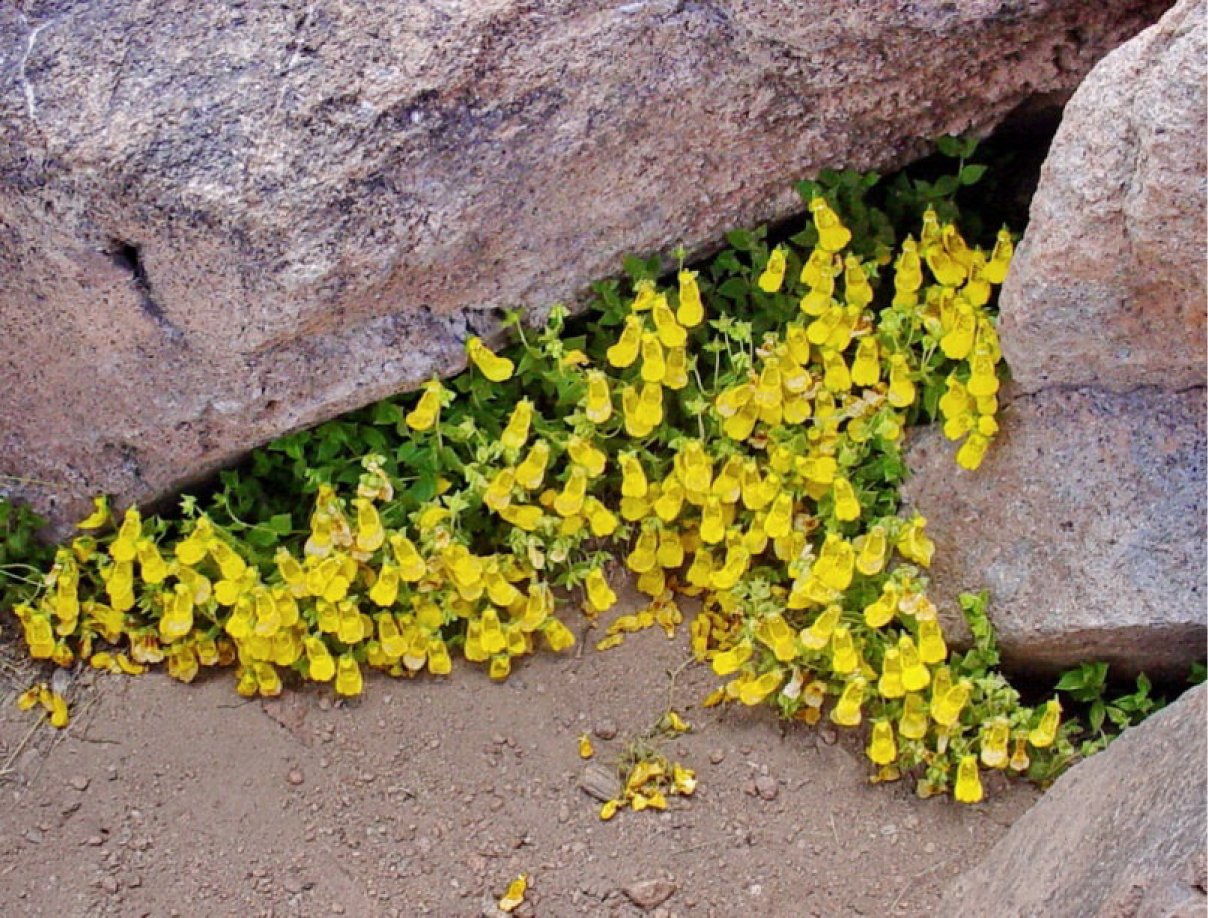
*Calceolaria stellariifolia*. (Photo - A.R. Flores, Fig. 6 Jan 2006)

**Fig. 9.**
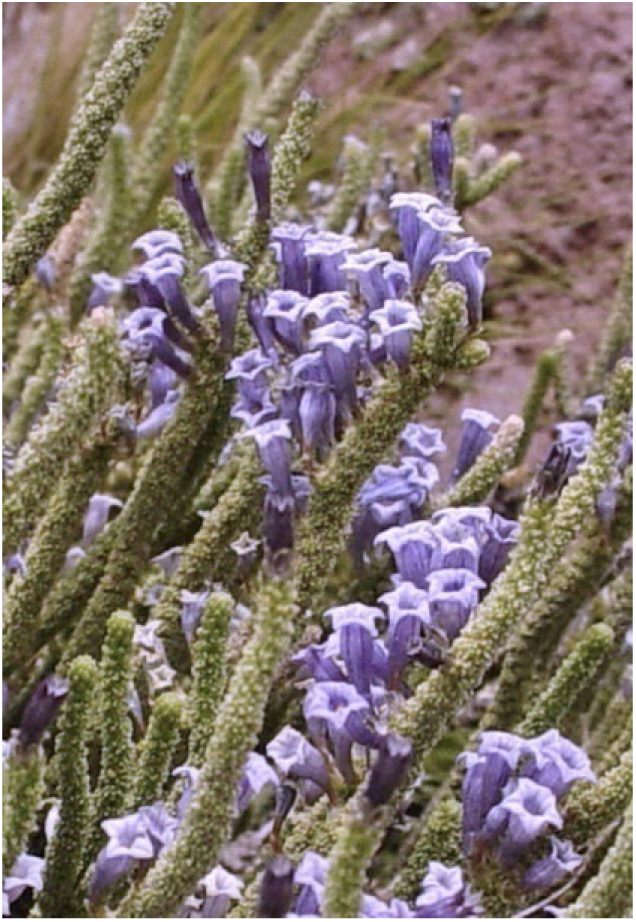
*Fabiana bryoides*. (Photo - A.R. Flores, 6 Jan 2006)

**Fig. 10.**
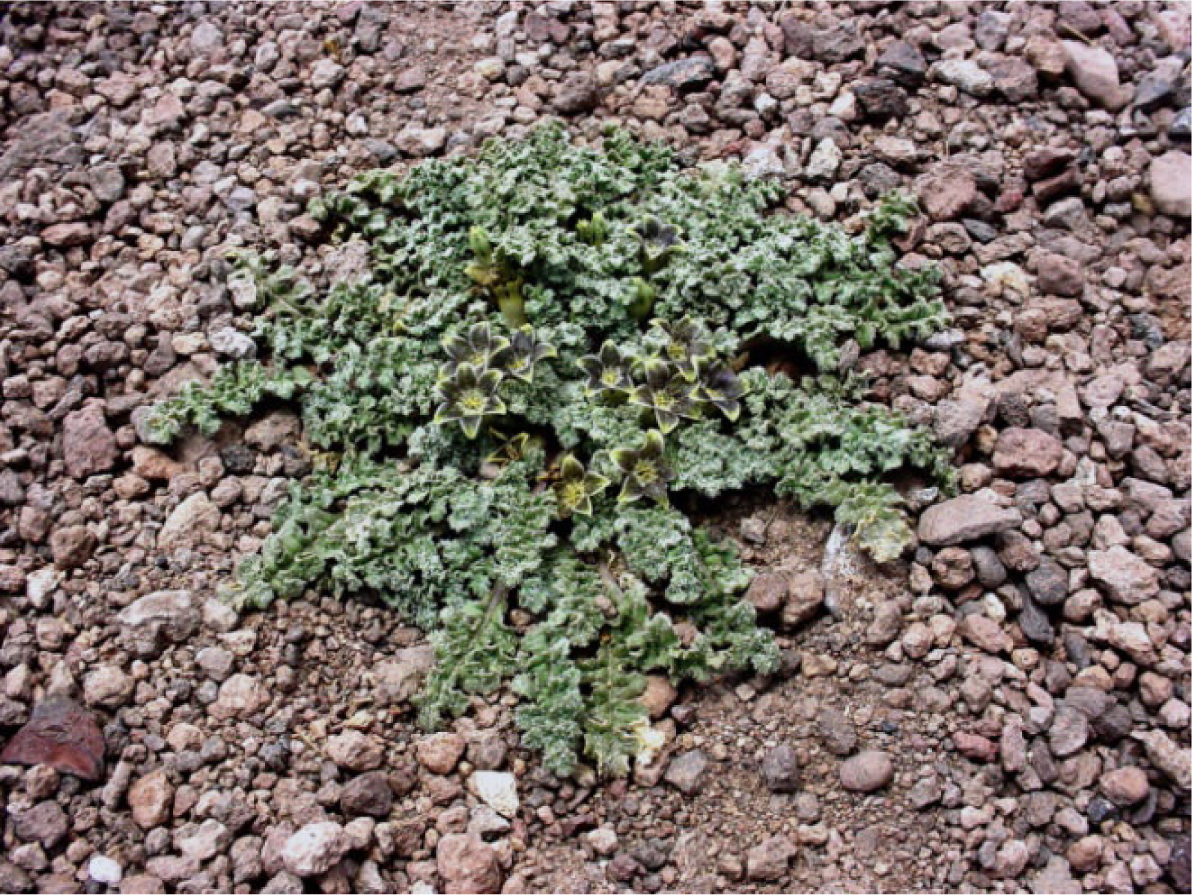
*Jaborosa parviflora*. (Photo - A.R. Flores, 6 Jan 2006)

**Fig. 11.**
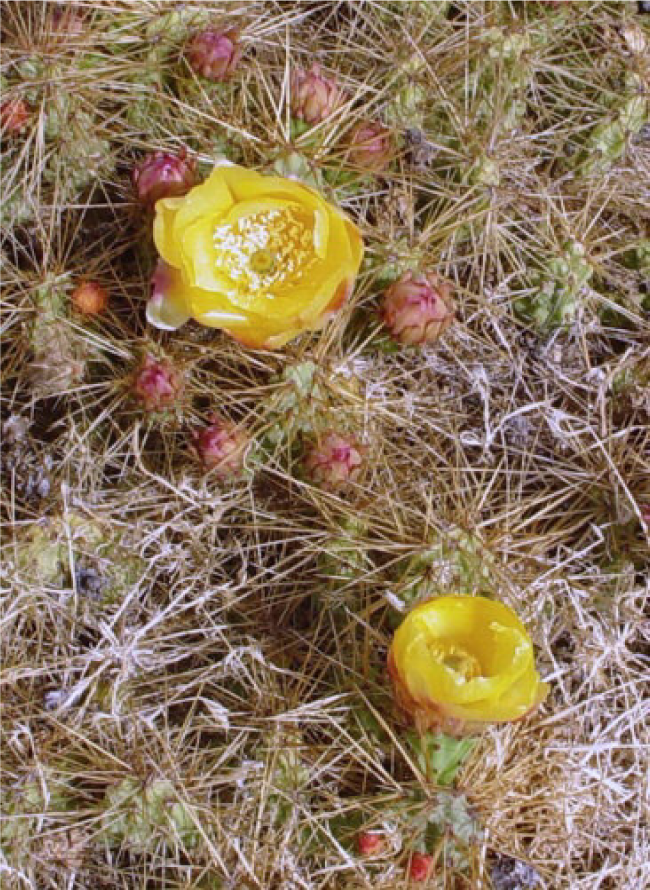
*Maihueniopsis boliviana*. (Photo - A.R. Flores, 6 Jan 2006)

#### Phenology

So far as can be judged, *V. unica* is presumed to flower in February, and seed dispersal may be assumed to follow upwards of a month after anthesis.

#### Etymology

*unica*, the Latin epithet we have provided for this new species, translates as ‘the only’, ‘the solitary’, which indicates our knowedge of it as no more than one individual plant with a single rosette. It also refers to its possession of a style crest with three strongly curved appendages, the upper opposed directionally to the lateral pair, which is without parallel to the present in the genus *Viola*.

#### Conservation assessment

Even although no information exists as to the size or extent of the only known population of the new species, with only one individual collected so far, the fact that it was found in an extensive area of open-cast mining operations leaves no doubt at all that it must qualify as EX (IUCN 2012): i.e. potentially in danger of extinction.

## Discussion: notes on species allied to *Viola unica* in section *Andinium*

A currently informal division of sect. *Andinium* results in a number of alliances, most of them very distinctive. Two are considerably more numerous than the rest, the largest being what we refer to as the broad *Viola volcanica* alliance, which itself splits naturally into smaller divisions. The relevant three subdivisions herein of this basic alliance are circumscribed by flexible, broad leaves; usually crenate margins which are more or less ciliate; and lamina faces with raised reticulate venation. Leaf glands may be present or not. Most species have no more than one apical style crest, although two species possess lateral lobes only, one on either side of the style head (Watson & Flores 2009, Watson *et al*. 2019a). The group concerned as whole extends from northern Patagonia (41°S) up to the border of the temperate and tropical zones (21°S).

A second much smaller alliance within sect. *Andinium* is known from central Andean to NW Argentinia. One of its published species has also recently been found in adjacent Chile but is not registered formally from there yet (Watson & Flores ined.). This group consists of four Argentinian endemic species described by Becker (1922, 1925, 1926, 1928), and one published earlier. The latter, *Viola flos-idae* (fig. 12), has not been recognised as sharing the triflabellate style crest until this present publication, despite having been drawn in detail in Rossow *et al*. (2003). As noted above, *V. mesadensis* may also belong here. The alliance, the *Triflabellatae*, is so far the only infrasectional grouping to be outlined and described in the literature. It was published without a taxonomic rank (Becker 1926), and is distributed between the provinces of Mendoza in the south (33°S) and Salta (25°S) to the north. The species were unambiguously defined by their style crests, consisting of two lateral lobes, one on either side of the style head, and one apical lobe. This configuration is shared by a few unrelated annual species of sect. *Andinium*, but for perennials of the genus it has hitherto been limited to these six species. *V. mesadensis* has fairly rounded laminas of a very similar outline to those of *V. unica*, and with the same number of marginal crenations, but almost twice as long and wide.

**Fig. 12.**
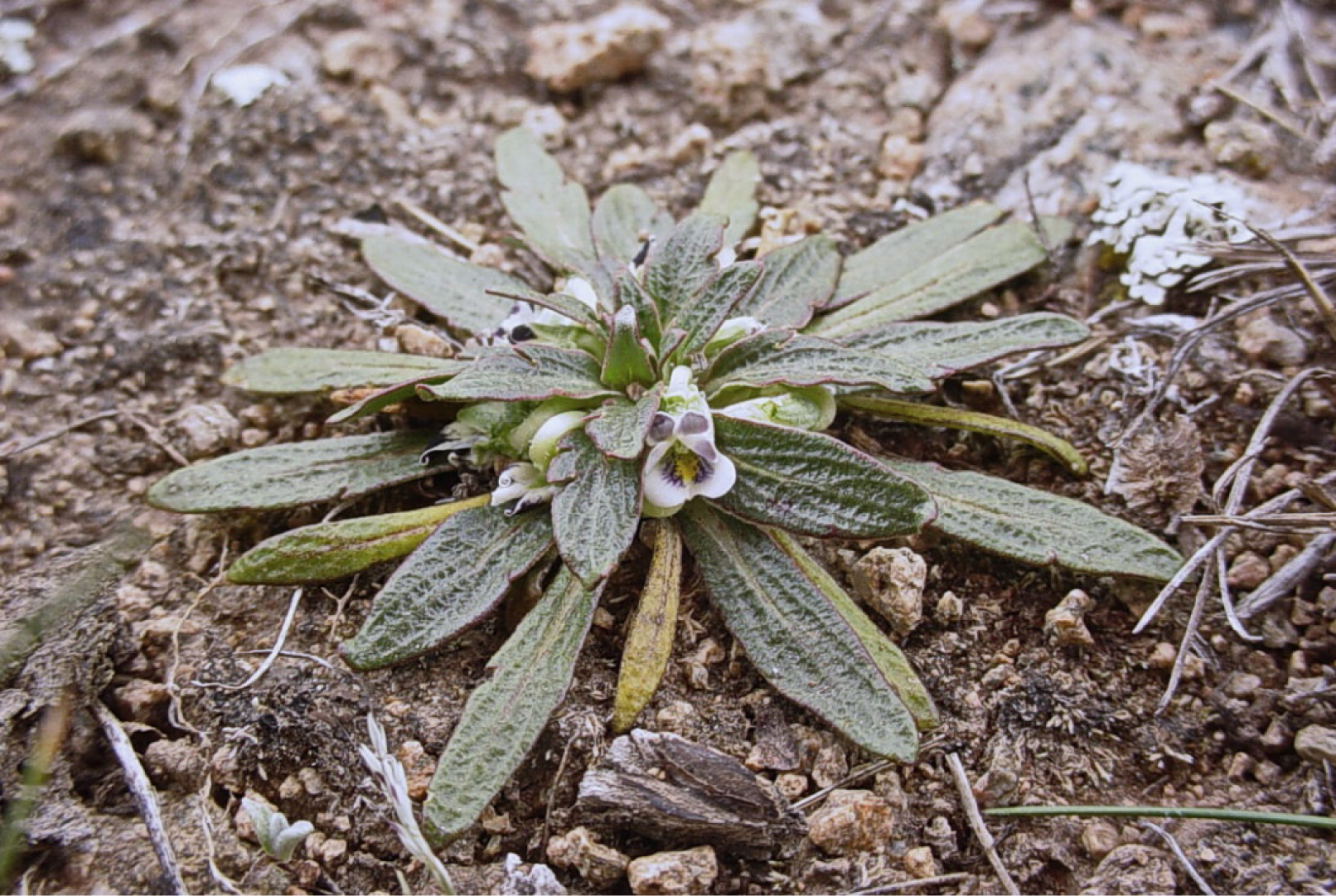
*Viola triflabellata*. (Photo - A.R. Flores, 14 Mar 2007)

As also do *V. flos-idae* and *Viola joergensenii* W. Becker (fig. 13), the new *V. unica* posesses distinct morphological characteristics of both the *V. volcanica* alliance (lamina morphology) (figs, 1-3) and the *Triflabellatae* (style crest) (fig. 4), and may thus be considered intermediate between the two groups. Although the possibility of it having a hybrid origin, or that the crest may have evolved independently, certainly cannot be ruled out, our assessment indicates that for all three species it is most likely an evolutionary step from the former alliance to the latter. As noted, sect. *Andinium* evolved in what is now the extreme south of the subcontinent (Clausen 1929, Ballard *et al*. 1999, Marcussen *et al*. 2012, Marcussen *et al*. 2015). This indicates that the general geographical direction of evolution of its taxa would have been from south to north, which supports the probability of it being intermediate rather than a natural hybrid. Two recently described species located within the *V. triflabellata* distribution range, *Viola beati* J.M. Watson & A.R. Flores (fig. 14) and *Viola singularis* J.M. Watson & A.R. Flores, are characterized by lateral lobes (Watson & Flores 2009, 2014, Watson *et al*. 2019a). Apart from *Viola llullaillacoensis* W. Becker (apical crest) (fig. 15), also a Chilean endemic of the high Altiplano, all others of the *V. volcanica* alliance *sensu stricto* lie to the south of the *Triflabellatae* range except for four rare and local species with lateral or apical crests only, *Viola exsul* J.M. Watson & A.R. Flores, *Viola gelida* J.M. Watson, M. Cárdenas & A.R. Flores (fig. 16), *Viola roigii* Rossow (fig. 17) and *Viola xanthopotamica* J.M. Watson & A.R. Flores (fig. 18). The only two taxa with an equivalent loose disposition of the rosette to that of *V. unica*, with its few and separated leaves, are *V. beati* and *V. singularis*, but those have a woody aerial framework: *V. unica* does not. The placement of *Viola mesadensis* W. Becker is unclear. It is only known from the type specimen, which was destroyed at B during WW2 (Haagemann & Zepernick 1993, Hiepko 1987), and our attempt in 2007 to rediscover it at the type location came to nothing. Becker (1928) was unsure whether it was either annual or perennial, the same problem facing us with *V. unica*. He described the style crest as consisting of three recurved laciniae, one from the apex, the other two lateral. This effectively is the broad equivalent of his *Triflabellatae*, although he did not place the species in that alliance, despite having published it himself two years earlier (Becker 1926). It is possible he considered the morphology of its laciniae to be too remote from the otherwise broad lobes of the *Triflabellatae*. Considering the equivalent of its foliage to that of the three lobed *V. flos-idae* and *V. joergensenii*, as well as its possible perennial life form, we consider it should provisionally be included in the *Triflabellatae*.

**Fig. 13.**
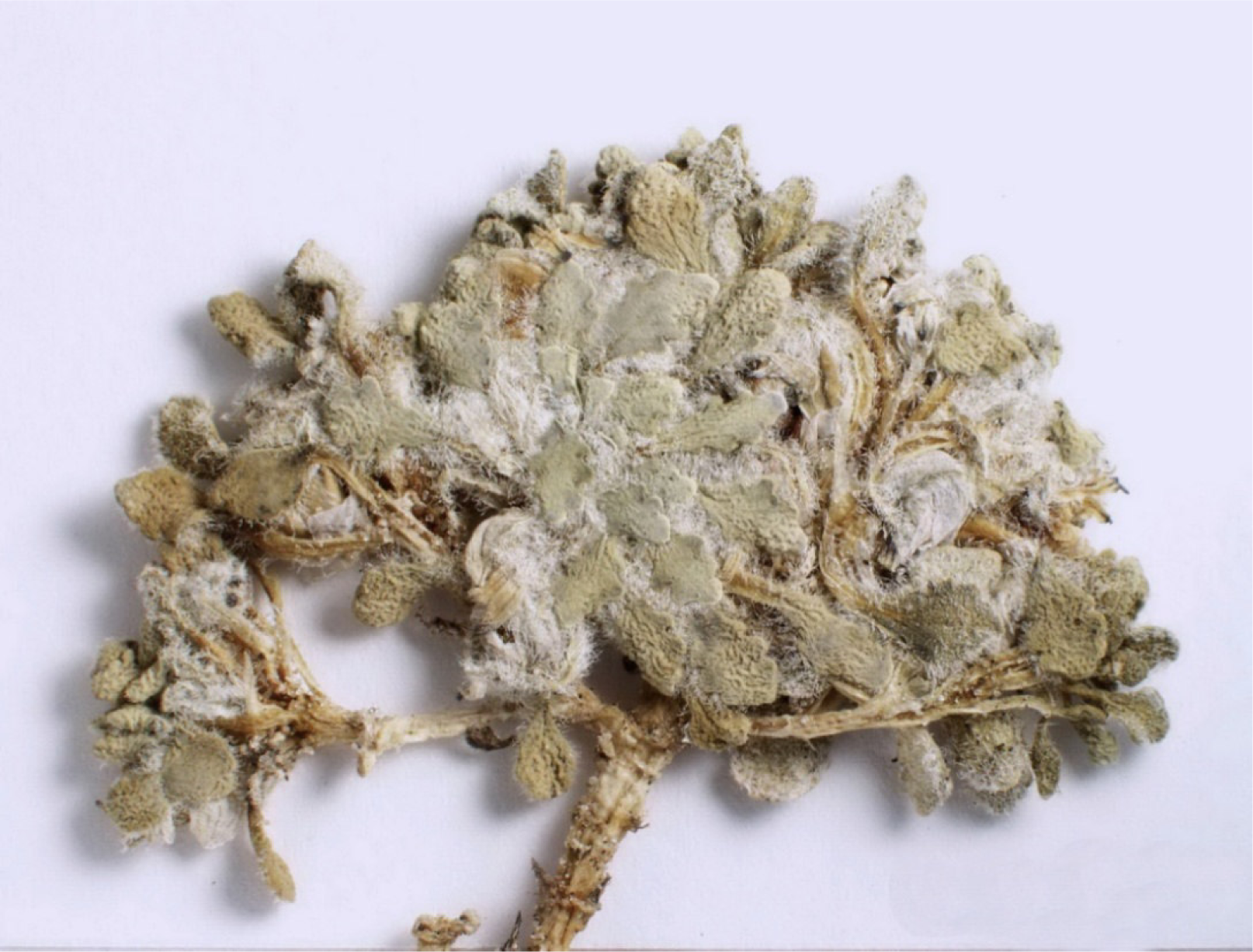
*Viola beati*. (Courtesy of BGBM and Fr. Monika Lüchow)

**Fig. 14.**
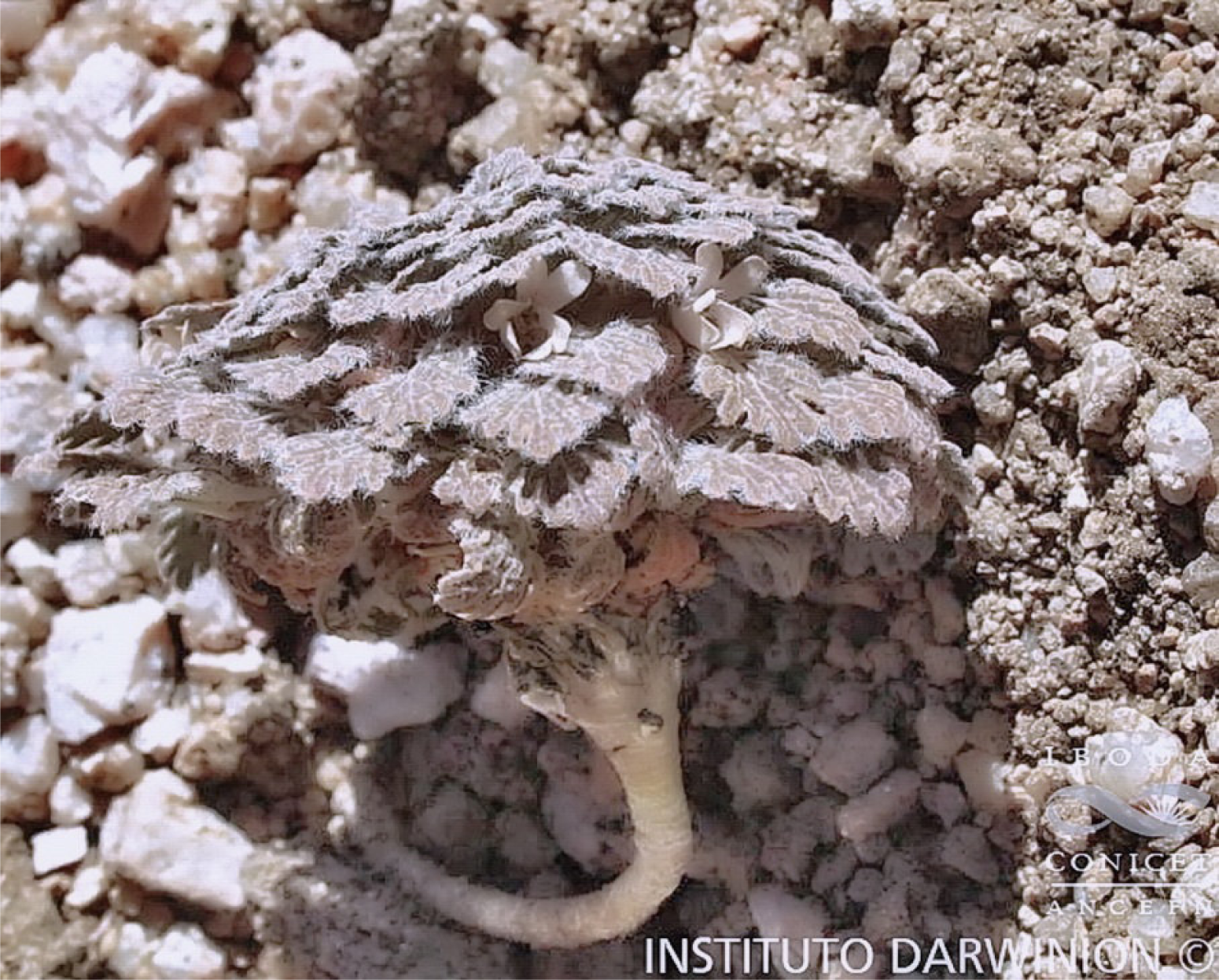
*Viola flos-idae*. (Photo - 28 Jan 2013. Courtesy of Instituto Darwinion, Buenos Aires)

**Fig. 15.**
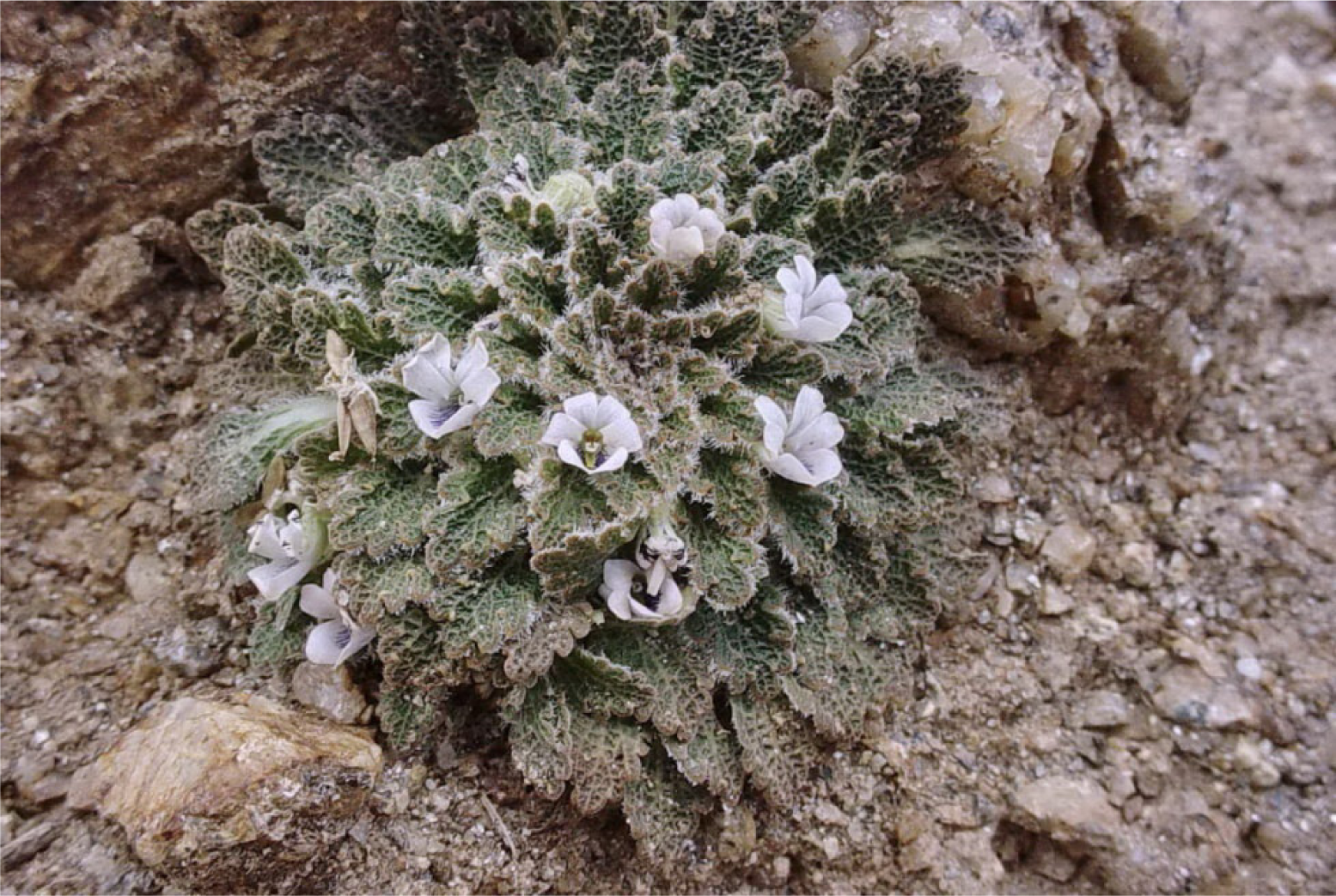
*Viola joergensenii*. (Photo - A.R. Flores, 13 Feb 2007)

**Fig. 16.**
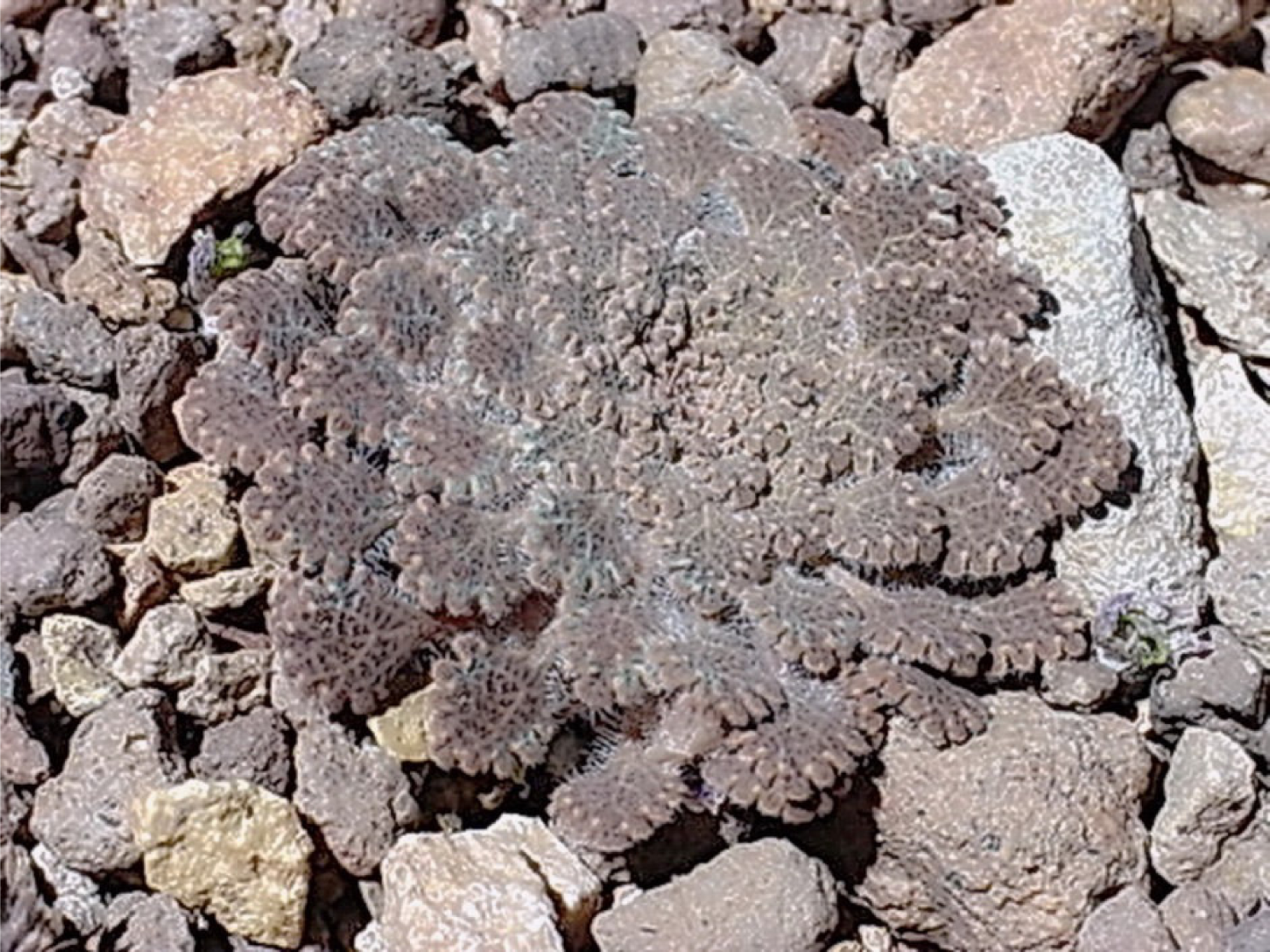
*Viola llullaillacoensis*. (Photo anon., ex Internet)

**Fig. 17.**
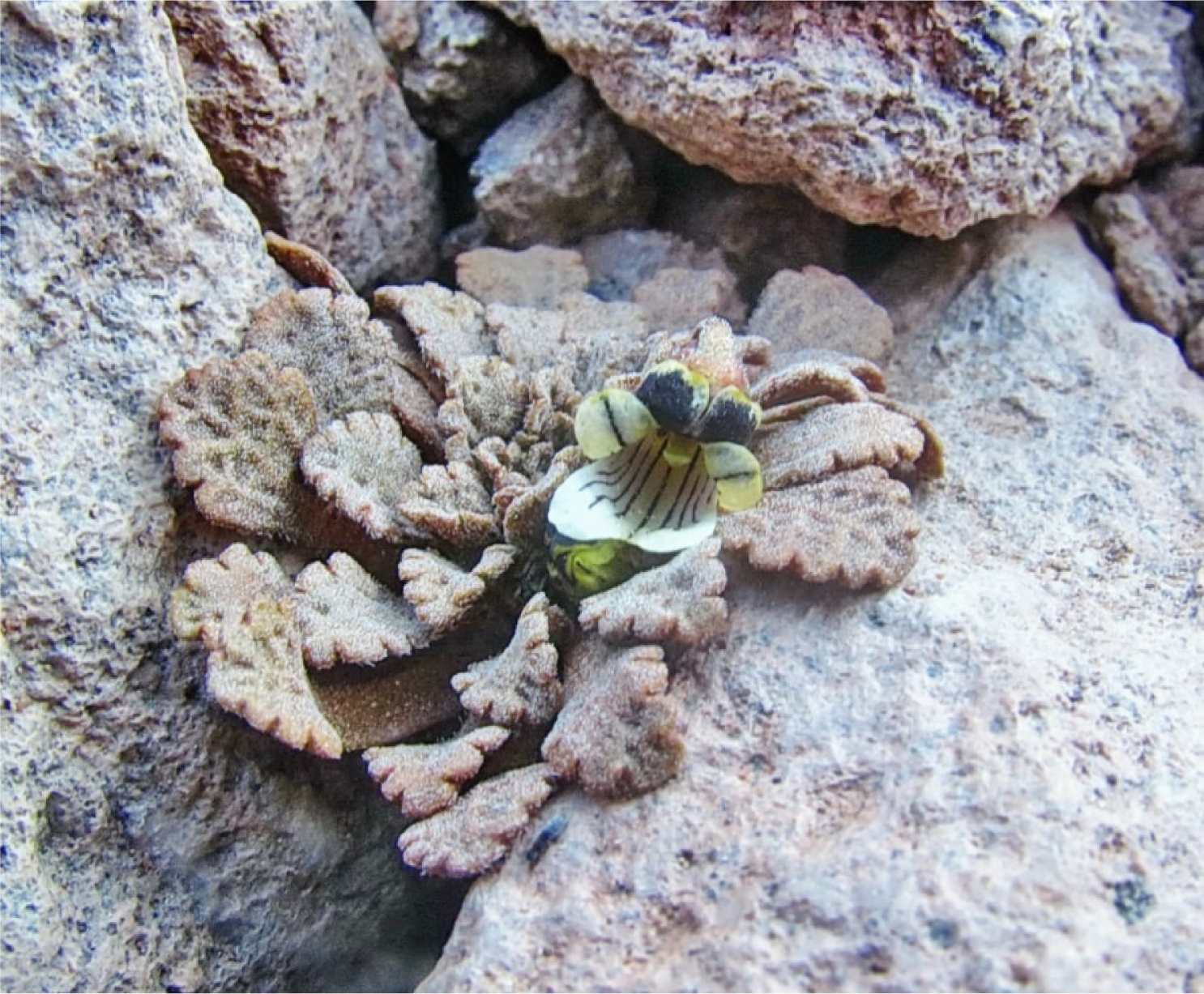
*Viola gelida*. (Photo - 11 Dec 2011. Courtesy of M.P. Cárdenas and Cedrem)

**Fig. 18.**
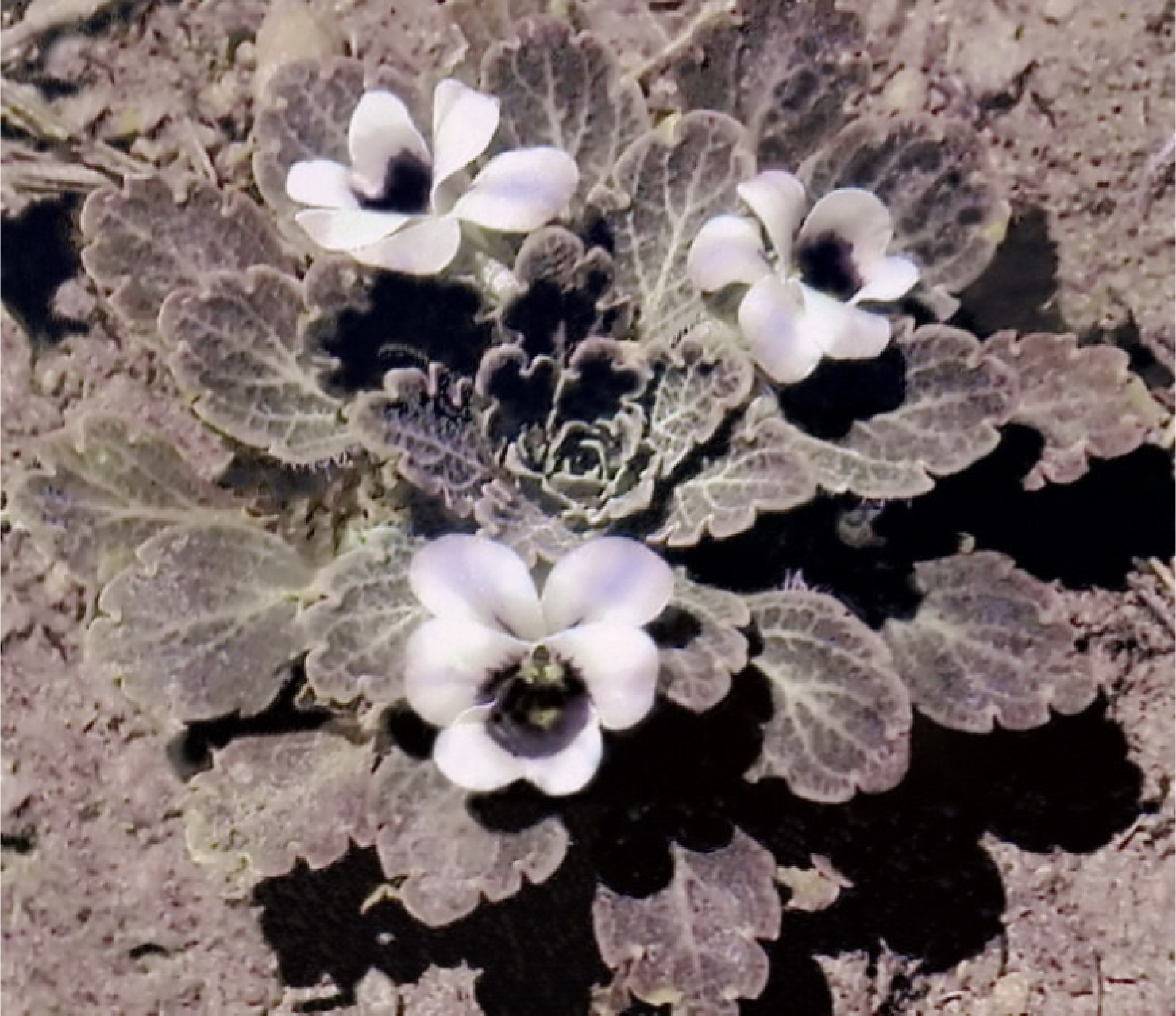
*Viola roigii*. (Photo - Roberto Kiesling)

Although further detailed investigation is required to determine the precise position of *V. unica* with respect to other species of sect. *Andinium* apparently related to it, we tentatively place it in the *Triflabellatae* as well. As such it is the second of that alliance to be recorded from Chile, and first endemic to that nation.

In order to clarify this situation as far as possible given present knowledge, we provide a key as follows to disinguish all species which may be significantly allied morphologically in any way to *V. unica*.

## Key to differentiate species of *Viola* sect. *Andinium* considered to be related to *Viola unica* which possess some combination of equivalent or relevant lamina face, lamina margin, crest morphology, and corolla coloration

1. Style crest apical or lateral … 2. - Style crest apical and lateral … 17.
2. Plant acaulous. Style crest apical … 3. - Plant with aerial ligneous structure. Style crest as lateral lobe on either side of style head … 16.
3. Lamina entire, as wide as long, or more … 4. - Lamina crenulate or subcrenulate … 5.
4. Margin distinctly red. (Endemic of N Argentinian Patagonia) … *Viola rubromarginata* - Margin concolorous with lamina face. (Endemic of N Argentinian Patagonia) … *Viola trochlearis*
5. Style crest entire, subrotund … 6. - Style crest entire, truncate or trilobed … 7.
6. Lamina face hyaline-setaceous. (Endemic of northern central Argentina) … *Viola exsul* - Lamina face minutely white papillose. (Endemic of northern Chile) … *Viola gelida*
7. Upper lamina surface pilose. (Endemic of northern Chilean Altiplano) … *Viola llullaillacoensis* - Upper lamina surface glabrous … 8.
8. Lamina with sinus glands at base of marginal crenations … 9. - Lamina without sinus glands at base of marginal crenations … 11.
9. Lamina undersurface usually without glands; or few when present. (Endemic of central southern Chile) … *Viola chillanensis* - Lamina undersurface always densely covered with glands … 10.
10. Lamina upper surface prominently alveolate reticulate, apex obtuse. (Central Argentina and central to central southern Chile) … *Viola congesta* - Venation of lamina upper surface not prominent, apex acute. (Central southern Chile and Argentinian northern Patagonia) … *Viola farkasiana*
11. Style crest entire, undersurface of lamina with glands … 12. - Style crest trilobed, Undersurface of lamina eglandular … 13.
12. Lamina margin incised-crenate. Calyx glabrous. (Argentina, from S Mendoza Province to northern Patagonia, and central southern Chile) … *Viola volcanica* - Lamina margin subentire. Calyx hirsute. (Central southern Chile and Argentinian northern Patagonia) … *Viola rugosa*
13. Lamina with 4 or more marginal crenations per side … 14. - Lamina with 3 marginal crenation per side … 15.
14. Lamina margin glabrous to few-ciliate at base. Lateral petals bearded. (Endemic of NW Argentina) … *Viola xanthopotamica* - Lamina margin distinctly ciliate. Lateral petals glabrous. (Endemic of central Chile) … *Viola friderici*
15. Marginal crenations of lamina large, tips flat. (Endemic of W central Argentina) … *Viola roigii* - Marginal crenations of lamina small, tips rounded. (Endemic of central Chile) … *Viola exilis*
16. Undersurface of lamina with glands. (Endemic of NW Argentina) … *Viola beati* - Undersurface of lamina eglandular. (Endemic of NW Argentina) … *Viola singularis*
17. Undersurface of lamina with evident dark, linear glands … 18. - Undersurface of lamina eglandular, or very rarely with few inconspicuous glands … 19.
18. Lateral petals glabrous. (Endemic of NW Argentina) … *Viola mesadensis* - Lateral petals bearded with clavate hairs. (Endemic of NW to W central Argentina and Altiplano of northern Chile) … *Viola flos-idae*
19. Lamina as wide as long, or wider, apex obtuse. (Endemic of northern Chilean Altiplano) … ***Viola unica*** - Lamina narrowly elliptical to ovate, apex acute or subacute … 20.
20. Margin of lamina glabrous … 21. - Margin of lamina ciliate … 22.
21. Stipules entire, 2-3 mm long. Leaf 1 cm long. (Endemic of NW Argentina) … *Viola triflabellata* - Stipules lacerate-fimbriate, 5-6 mm long. Leaf 2-3 cm long. (Endemic of NW Argentina) … *Viola hieronymi*
22. Stipules entire. Style crest lobes sessile. (Endemic of NW Argentina) … *Viola joergensenii* - Stipules fimbriate. Style crest lobes stipitate. (Endemic of NW Argentina) … *Viola tucumanensis*

The same taxa listed as they occur naturally from south to north, in each case based on the closest known population to *V. unica*. Species of the *Triflabellatae* published previously and recognised herein are printed in bold type. Two species tentatively assigned to that alliance, including the novelty presented here, are underlined.

*Viola trochlearis* J.M. Watson & A.R. Flores (fig. 19)

**Fig. 19.**
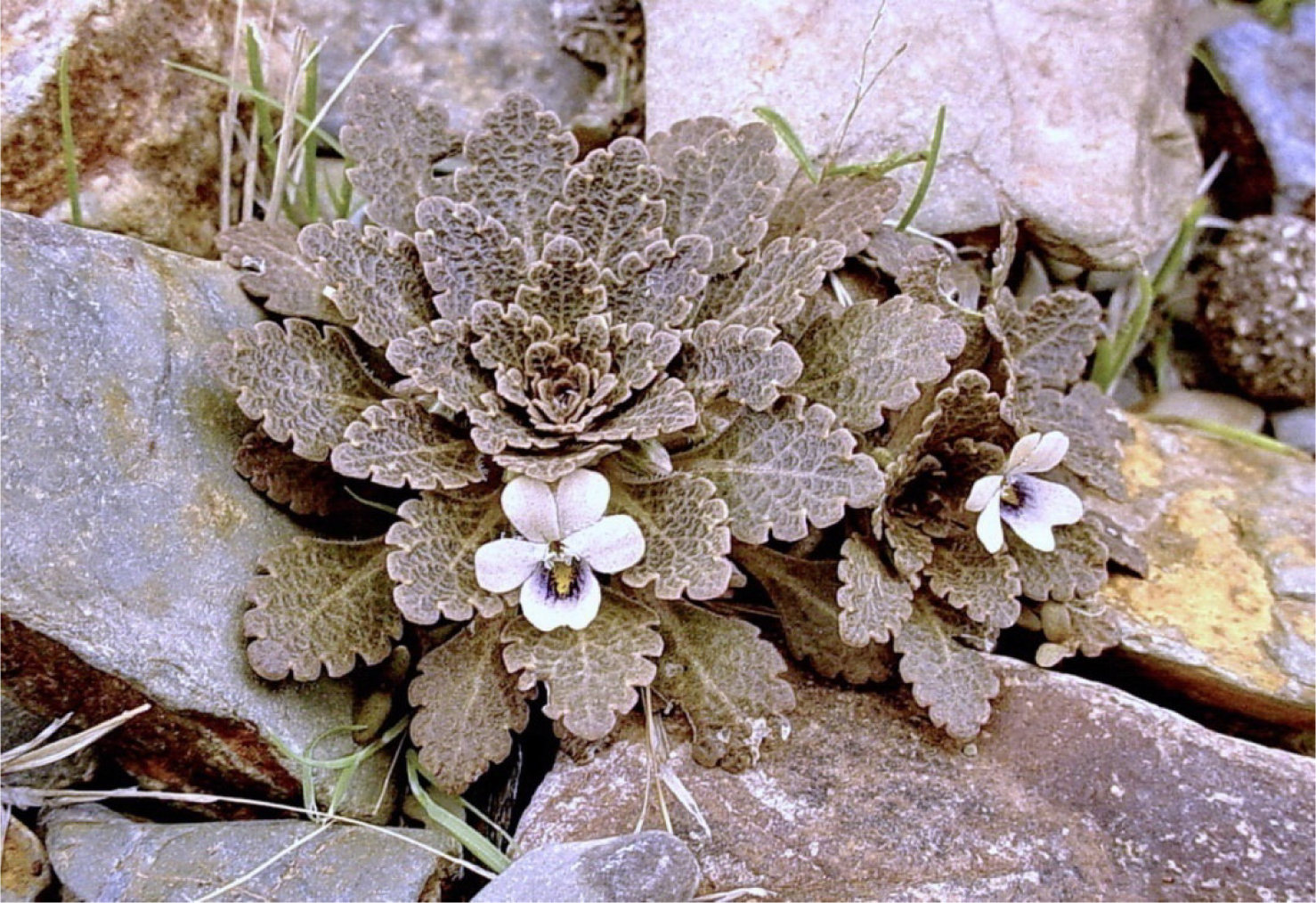
*Viola xanthopotamica*. (Photo - A.R. Flores, 9 Feb 2007)

*Viola rugosa* Phil. ex W. Becker (fig. 20)

**Fig. 20.**
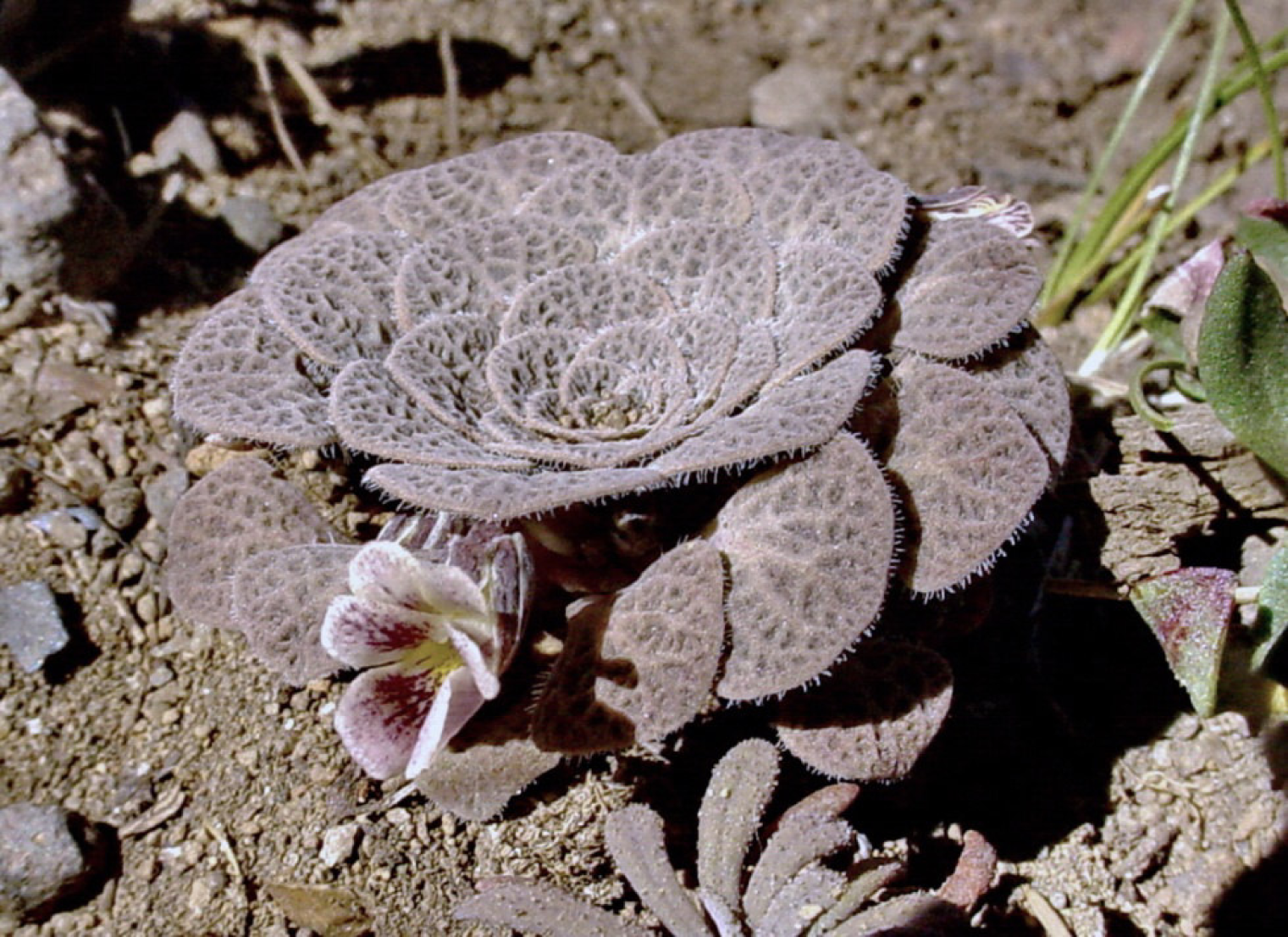
*Viola trochlearis*.(Photo - A.R. Flores, 21 Dec 2007)

*Viola chillanensis* Phil. ex W. Becker

*Viola farkasiana* J.M. Watson & A.R. Flores (fig. 21)

**Fig. 21.**
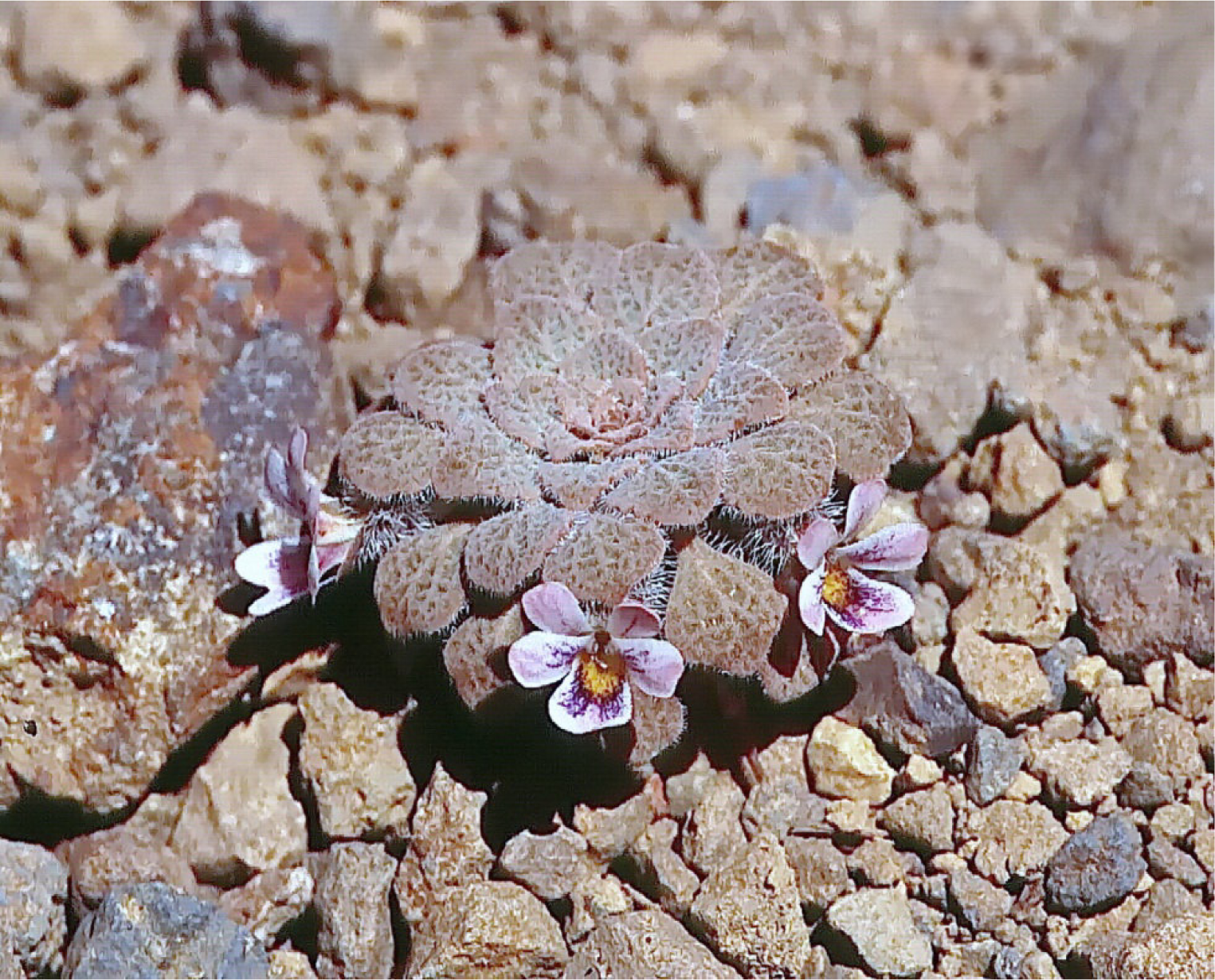
*Viola rugosa*. (Photo - J.M. Watson, 7 Feb 2003)

*Viola rubromarginata* J.M. Watson & A.R. Flores (fig. 22)

**Fig. 22.**
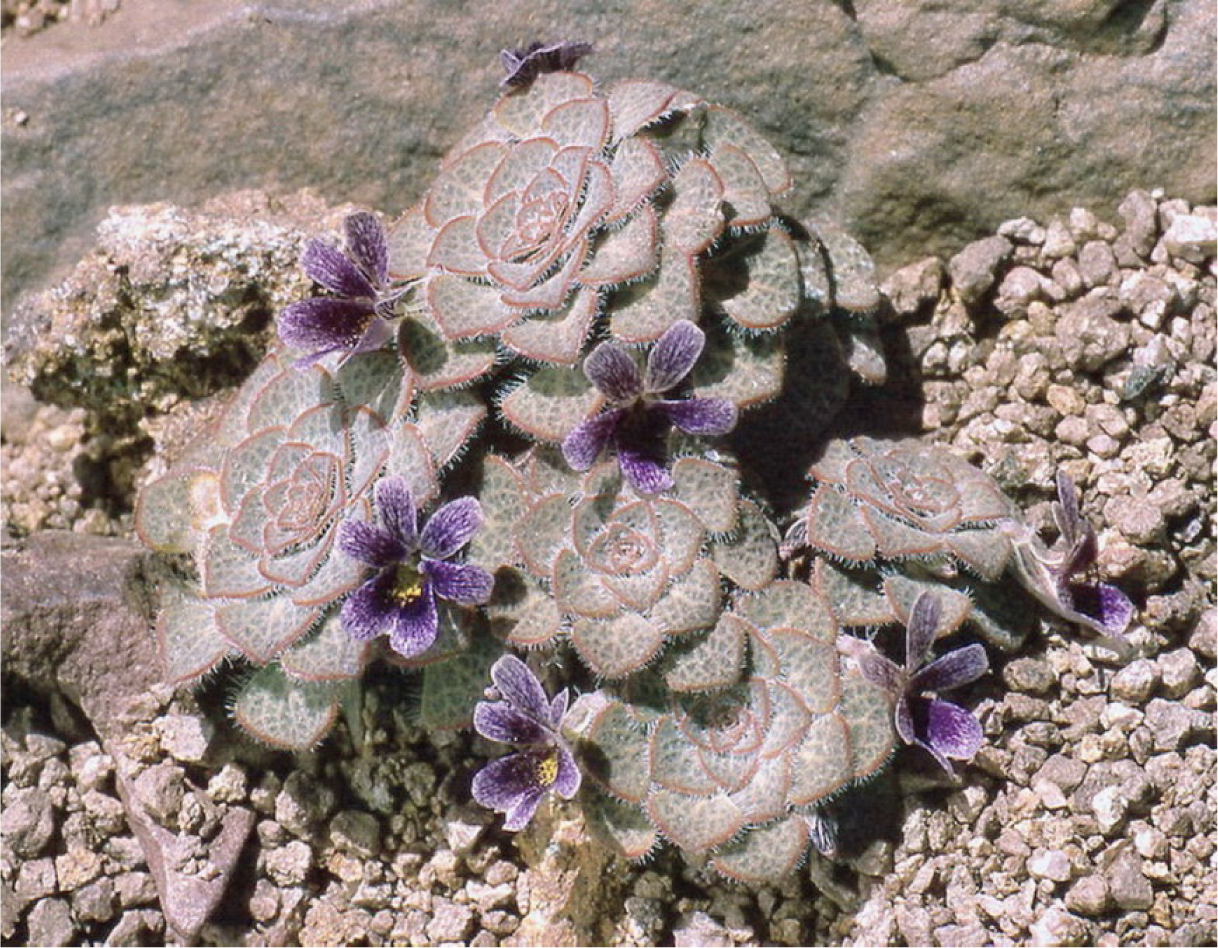
*Viola rubromarginata*. (Photo - J.M. Watson, Dec 1994)

*Viola volcanica* Gillies ex Hook. & Arn. (fig. 23)

**Fig. 23.**
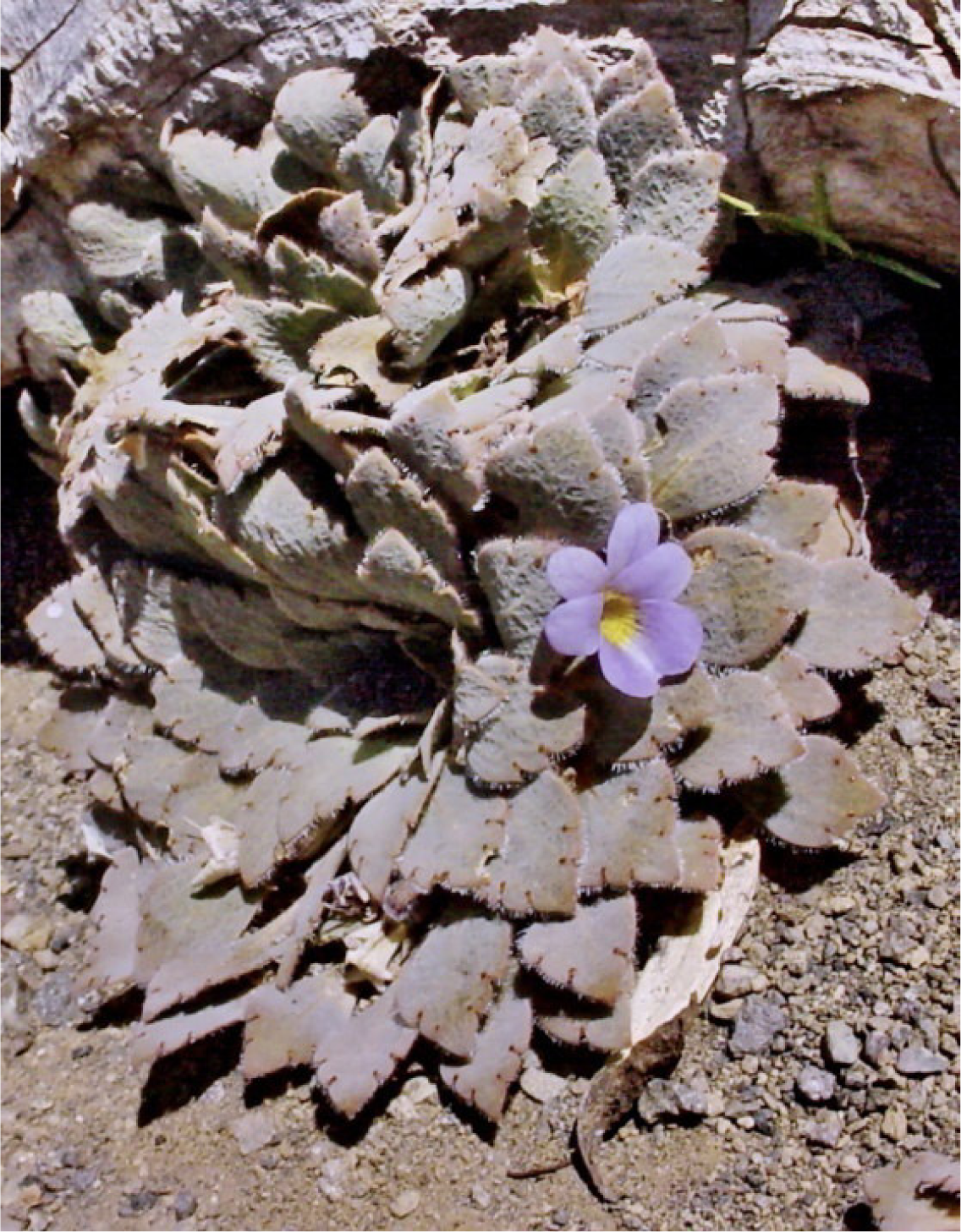
*Viola farkasiana*. (Photo - A.R. Flores, 25 Dec 2002)

*Viola congesta* Gillies ex Hook. & Arn. (fig. 24)

**Fig. 24.**
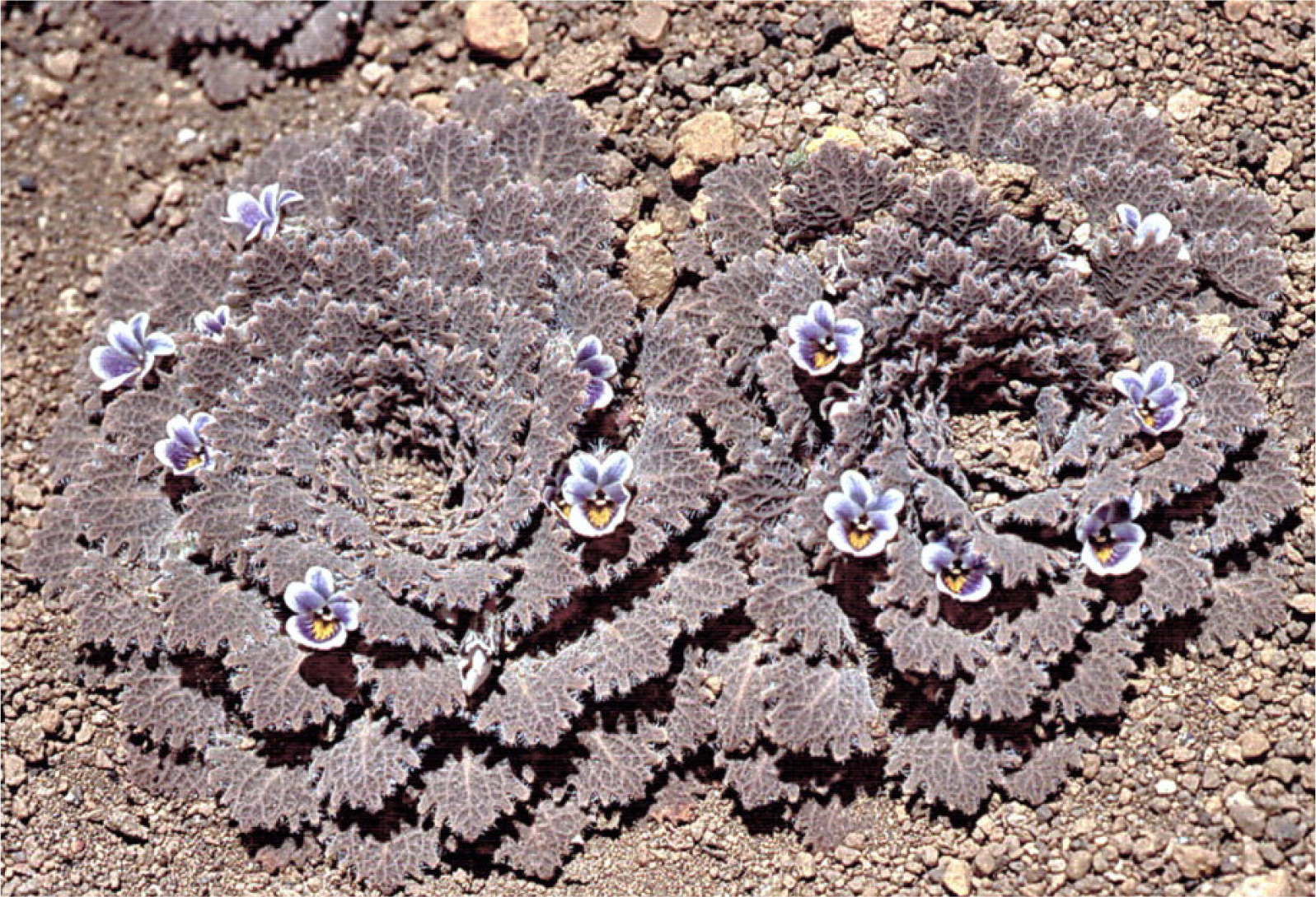
*Viola volcanica*. (Photo - J.M. Watson, 29 Dec 2002)

*Viola friderici* W. Becker

*Viola exilis* Phil.

*Viola roigii* Rossow (fig. 17)

*Viola exsul* J.M. Watson & A.R. Flores

*Viola gelida* J.M. Watson, M.P. Cárdenas & A.R. Flores (fig. 16)

***Viola flos-idae*** Hieron. (fig. 12)

*<u>Viola mesadensis</u>* W. Becker

*Viola xanthopotamica* J.M. Watson & A.R. Flores (fig. 18)

***Viola hieronymi*** W. Becker

***Viola joergensenii*** W. Becker (fig. 13)

*Viola beati* J.M. Watson & A.R. Flores (fig. 14)

*Viola singularis* J.M. Watson & A.R. Flores

***Viola triflabellata*** W. Becker (fig. 25)

**Fig. 25.**
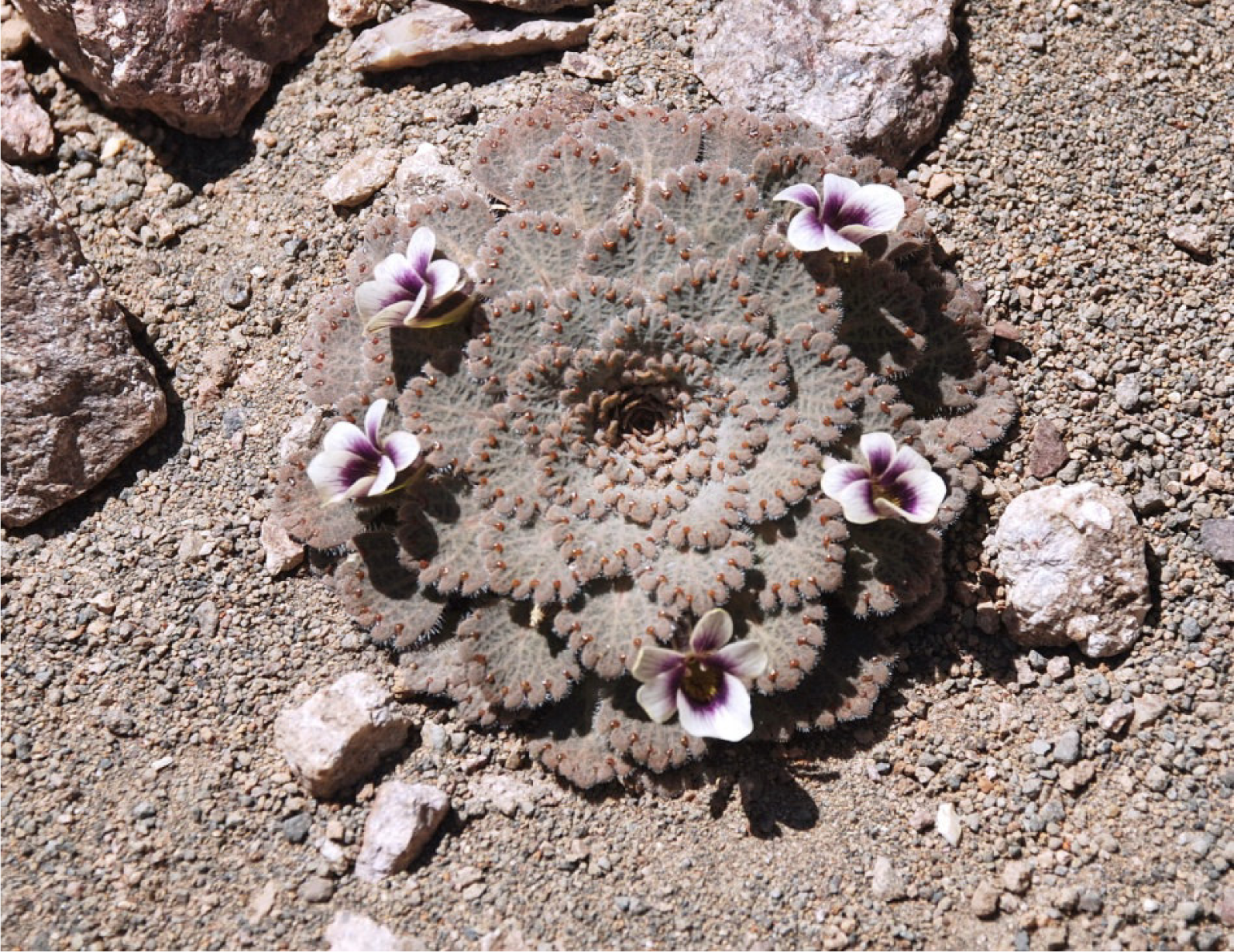
*Viola congesta*. (Photo - J.M. Watson, 17 Dec 2013)

***Viola tucumanensis*** W. Becker

*Viola llullaillacoensis* W. Becker (fig. 15)

*<u>Viola unica</u>* J.M. Watson & A.R. Flores (figs. 1-4)

### Note 4

In her treatment of the violas listed for Argentina, Nicola (2017) synonymises *V. joergensenii* and *V. tucumanensis* with *V. triflabellata*, describing the latter as an extremely polymorphic species. Photographs of the first and last of those taxa taken in the field by ourselves show that they are distinct beyond dispute.

Becker gives precise stipule lengths for each of his species: 10 mm only for *V. joergensenii*; 5 mm only for *V. tucumanensis*; 2-3 mm for *V. triflabellata*. The question then arises - Why is 5 mm given as the minimum length for *V. triflabellata* by Nicola (with 10 mm as the maximum)? If we take Becker’s measurements into account, as we certainly should, then the stipules of *V. triflabellata* as recognised by Nicola vary between 2-10 mm in length, and are either glabrous or ciliate. This strikes us as frankly absurd in the context of the section. To draw from Nicola (2017) herself, measurements of the stipules given for the 14 cited Argentinian taxa of the section which possess them, and excluding *V. triflabellata*, range from 1-6 mm in length. The average longitudinal variation noted for each is 0.8 mm, with a rare maximum of 2.5 mm. That compares with hers and Becker’s combined 8 mm for *V. triflabellata*.

We have photographed and gathered specimens in the immediate general sector of the *V. tucumanensis* type location in Salta Province which do not correspond to Becker’s protologue of that taxon. They are, however, identical in almost every respect to a population of *V. triflabellata* recorded by Becker and registered by us in La Rioja Province, over 400 km to the south. This suggests that *V. triflabellata* is in fact very morphologically stable, and occurs in the same vicinity as *V. tucumanensis* in Salta.

Given these facts, we will maintain Becker’s original judgement unless clear, unambiguous evidence to the contrary can be produced.

Of three sympatric perennial species with apical and laterally lobed style crests included in her entry (Nicola 2017), *V. hieronymi* and her concept of *V. trifabellata* are discussed and differentiated. However, this morphology is also present in *V. flos-idae*, a factor she overlooked and failed to consider in a wider context, even though the accompanying drawing and her description present the form of the crest unambiguously. Along with at least one other legitimately published and recognised species, trilaciniate *V. mesadensis* is absent from Nicola’s treatment.

### Note 5

*Viola flos-idae* is polymorphic in leaf shape and number of marginal crenations, including rarely to almost entire. The lateral petals may possess either clavate hairs or a basal tumour. The apical lobe of the style crest is sometimes itself bilobed. Importantly, very occasionally the underleaf glands are absent on a few leaves (Rossow *et al*. 2003). Overall *V. joergensenii* shares most of the morphological features of *V. flos-idae*, including a markedly reticulated upper lamina face with regular small, rounded crenations on the margins (pers. obs.), corolla size, and basal indumentum on the inferior petal. In his protologue of *V. joergensenii*, Becker (1926) indicated that the reverse of the lamina has a few small glands. However, all plants we have examined from an extensive population near the type site are completely eglandular. In addition the two taxa are noted as sympatric (e.g. Rossow *et al*. 2003), although this needs informed confirmation. Whether *V. joergensenii* should be included as a synonym of *V. flos-idae* requires further careful investigation in view of their extreme superficial similarity, but at present we maintain them as distinct taxa on the basis of the following *V. flos-idae* vs *V. joergensenii* characters:

1. Lamina to as wide as long vs always ca. twice as long as wide.
2. Lamina margin with up to 9 crenations vs up to 4 crenations.
3. Lamina undersurface almost always glandular vs almost always eglandular.
4. Sepals usually glandular vs always eglandular.
5. Lateral petals with clavate hairs or basal tumours vs always with clavate hairs only.

*Viola flos-idae* is the most southerly and widespread of these triflabellate taxa. Considering the deduced geographical advance and linked evolution by sect. *Andinium* (Watson & Flores 2012, 2013a, b), as also covered in more detail above, it is probably the most ancestral of them as well. Without doubt, these two species and *V. mesadensis* are morphologically the closest of the *Triflabellatae* to *V. unica*.

## Acknowledgement

We are most grateful to the staff of the Santiago National Natural History Museum herbarium (SGO) for permission to visit and for helpful assistance.

## References

Ballard, H.E., jr., Sytsma, K.J. & Kowal, R.R. (1999) Shrinking the violets: Phylogenetic relationships of infrageneric groups in *Viola* (Violaceae) based on internal transcribed spacer DNA sequences. Syst. Bot. 23: 439–458.

Becker, W. (1922) Violae novae Americae meridionalis. Repert. Spec. Nov. Regni Veg. 18: 182, 184, 185.

Becker, W. (1925) Beiträge zur Kenntnis der südamerikanischen Violae. Repert. Spec. Nov. Regni Veg. 21: 351, 352, 357, 358.

Becker, W. (1926) Beiträge zur Violenflora Argentiniens. Repert. Spec. Nov. Regni Veg. 22: 350–354.

Becker, W. (1928) Ein Beitrag zur Violenflora Argentiniens und Chiles. Repert. Spec. Nov. Regni Veg. 24: 363–365.

Clausen, J. (1929) Chromosome number and relationships of some North American species of Viola. Ann. Bot. 63: 741–764.

Gajardo, R. (1994) La vegetación natural de Chile: classificación y distribución geográfica. Editorial Universitaria, Santiago de Chile. 165 pp.

Haagemann, I. & Zepernick, B. (1993) On the history of the Berlin Botanic Garden. In Haagemann, I. & Zepernick, B. (eds.) (transl. Smith, L.) The Berlin-Dahlem Botanic Garden: 9–11. Förderkriess der naturwissenschaftlichen Museen Berlins e. V.

Hiepko, P. (1987) The collections of the Botanical Museum Berlin-Dahlem (B) and their history. Englera 7: 219–252.

IPNI (2019) The International Plant Names Index Published on the Internet. http://www.ipni.org (accessed 2 September 2019)

IUCN (2012) International Union for Conservation of Nature red list categories and criteria: Version 3.1. Second edition.

Marcussen, T., Heier, L., Brysting, A.K., Oxelman, B. & Jakobsen, K.S. (2015) From gene trees to a dated alloploidy network: insights from the angiosperm genus *Viola* (Violaceae). Syst. Biol. 64: 84–101.

Marcussen, T., Jakobsen, K.S., Danihelka, J., Ballard, H.E., jr., Blaxland, K., Brysting, A.K. & Oxelman, B. (2012) Inferring species networks from gene trees in high polyploid North American and Hawaiian violets (*Viola*, Violaceae). Syst. Biol. 61: 107–126.

Nicola, M.V. (2017) Viola. In: Anton, A.M.R. & Zuloaga, F.O. (eds.) Flora vascular de la República Argentina 17: 371–408. Instituto Nacional de Tecnología Agropecuaria (INTA), Buenos Aires.

Rossow, R.A., Watson, J.M. & Flores, A.R. (2003) Violaceae. In: Kiesling, R. (ed.) Fl. San Juan 2: 142. Estudio Sigma, Buenos Aires.

Wahlert, G.A., Marcussen, T., de Paula-Souza, J., Feng, M. & Ballard, H.E., jr. (2014) A phylogeny of the Violaceae (Malpighiales) inferred from plastid DNA sequences: implications for generic diversity and intrafamilial taxonomy. Syst. Bot. 39: 239–252.

Watson, J.M. & Flores, A.R. (2009) A new and rare rosulate species of *Viola* from Argentina. Phytotaxa 2: 17–23.

Watson, J.M. & Flores, A.R. (2012) Fire and ice: rosulate viola evolution. Part one—the stage is set. Rock Gard. Quart., Bull. N. Am. Rock Gard. Soc. 70(4): 360–366.

Watson, J.M. & Flores, A.R. (2013a) Fire and ice: rosulate viola evolution. Part two—the drama unfolds. Rock Gard. Quart., Bull. N. Am. Rock Gard. Soc. 71(1): 42–53.

Watson, J.M. & Flores, A.R. (2013b) Fire and ice: rosulate viola evolution. Part three—a merry life and a short one. Rock Gard. Quart., Bull. N. Am. Rock Gard. Soc. 71(2): 118–141.

Watson, J.M. & Flores, A.R. (2014) Upping their number, assessing their risk. *Viola singularis* (Violaceae) revisited, and an evaluation of sect. *Andinium*, its higher taxonomic group. Phytotaxa 177(3): 177–182.

Watson, J.M. & Flores, A.R. (2018b) Triple alliance: new and rediscovered species of *Viola* section *Andinium* from Argentina. Int. Rock Gard. 106: 2-33. Published on the Internet: http://www.srgc.org.uk/logs/logdir/2018Oct251540501603IRG106October2018.pdf

Watson, J.M. & Flores, A.R. (2019a) Those damned promiscuous rosulates. A second new wild hybrid for *Viola* L. section *Andinium* W. Becker, also an Argentinian endemic. Int. Rock Gard. 113: 3-50. Published on the Internet: http://www.srgc.org.uk/logs/logdir/2019May311559296345IRG113.pdf

Watson, J.M. & Flores, A.R. (2019b) Follow the yellow quick flow. Another new species of *Viola* sect. *Andinium* from Argentina, this one from the country’s mountainous northwest. Int. Rock Gard. 115: 3-53. Published on the Internet: http://www.srgc.org.uk/logs/logdir/2019July251564083758IRG115.pdf

Watson, J.M., Flores, A.R. & Arroyo-Leuenberger, S. (2019) A new *Viola* (Violaceae) from the Argentinian Andes. Willdenowia 49(1); 35–41. https.//doi.org/10.3372/wi.49.49101

